# Biochemical and neurophysiological effects of deficiency of the mitochondrial import protein TIMM50

**DOI:** 10.1101/2024.05.20.594480

**Authors:** Eyal Paz, Sahil Jain, Irit Gottfried, Orna Staretz-Chacham, Muhammad Mahajnah, Pritha Bagchi, Nicholas T. Seyfried, Uri Ashery, Abdussalam Azem

## Abstract

TIMM50, an essential TIM23 complex subunit, is suggested to facilitate the import of ∼60% of the mitochondrial proteome. In this study, we characterized a *TIMM50* disease causing mutation in human fibroblasts and noted significant decreases in TIM23 core protein levels (TIMM50, TIMM17A/B, and TIMM23). Strikingly, TIMM50 deficiency had no impact on the steady state levels of most of its putative substrates, suggesting that even low levels of a functional TIM23 complex are sufficient to maintain the majority of TIM23 complex-dependent mitochondrial proteome. As TIMM50 mutations have been linked to severe neurological phenotypes, we aimed to characterize TIMM50 defects in manipulated mammalian neurons. TIMM50 knockdown in mouse neurons had a minor effect on the steady state level of most of the mitochondrial proteome, supporting the results observed in patient fibroblasts. Amongst the few affected TIM23 substrates, a decrease in the steady state level of components of the intricate oxidative phosphorylation and mitochondrial ribosome complexes was evident. This led to declined respiration rates in fibroblasts and neurons, reduced cellular ATP levels and defective mitochondrial trafficking in neuronal processes, possibly contributing to the developmental defects observed in patients with TIMM50 disease. Finally, increased electrical activity was observed in TIMM50 deficient mice neuronal cells, which correlated with reduced levels of KCNJ10 and KCNA2 plasma membrane potassium channels, likely underlying the patients’ epileptic phenotype.

## Introduction

The mitochondrion is a vital organelle found in nearly all eukaryotic cells, where it is involved in numerous important cellular functions and metabolic pathways, including supplying cellular energy, assembling iron-sulphur clusters, and regulating the cell cycle, cell growth and differentiation, programmed cell death, and synaptic transmission (1–3). In humans, these functions are executed by ∼1,500 different mitochondrial proteins, of which only 13 are encoded by the mitochondrial genome (4). The remaining mitochondrial proteins are nuclear-encoded and thus imported into the mitochondria. The translocated proteins are sorted to their specific mitochondrial compartment via several intricate protein translocation pathways, including the presequence pathway, which is used for import of nearly ∼60% of the mitochondrial proteins (5,6).

The TIM23 complex mediates the import of some intermembrane space (IMS) proteins, many mitochondrial inner membrane (MIM) proteins, and all mitochondrial matrix proteins (7). The TIM23 complex in yeast comprises three essential subunits, Tim23, Tim17 and Tim50 (TIMM23, TIMM17A/B and TIMM50 in mammals). Association of Tim21 (TIMM21 in mammals) and Mgr2 (ROMO1 in mammals) promotes the lateral translocation of proteins into the MIM, while association of the presequence translocase-associated motor (PAM) complex with the TIM23 core promotes the import of matrix proteins (8–10). Recent structural analysis showed that Tim17 forms the protein translocation path, whereas the associated Tim23 protein likely plays a structural role, serving as a platform that mediates the association of other complex subunits (11,12).

Tim50, first discovered in yeast some two decades ago (13,14), is thought to be the first TIM23 complex component to interact with presequences of precursor proteins as they emerge from the Tom40 channel, thus playing a pivotal role in presequence-containing protein sorting (15). It was further suggested that normal Tim50 functionality is required for maintaining the mitochondrial membrane potential (16). Additionally, TIMM50, the mammalian homologue of Tim50, was shown to be involved in steroidogenesis and plays a preventive role in pathological cardiac hypertrophy and several types of cancer (17–21).

Recently, TIMM50 has generated immense interest in human health research, as mutations in the encoding gene have been linked in geographically and ethnically varied populations to the development of a severe disease characterized by mitochondrial epileptic encephalopathy, developmental delay, optic atrophy, cardiomyopathy, and 3-methylglutaconic aciduria. To date, seven different mutations have been identified in children from ten unrelated families (22–27).

Although TIMM50 mutants are mostly associated with neurological disorders, functional characterization of TIMM50 has yet to be reported in neuronal cells. Additionally, despite being involved in the import of nearly 60% of the mitochondrial proteins, the impact of TIMM50 deficiency on the entire mitochondrial proteome has yet to be characterized. In this study, using a proteomics approach, we show the impact of TIMM50 deficiency on the mitochondrial and cellular proteome and characterize, for the first time, the neurological role of TIMM50 by knocking it down in mouse neurons.

## Results

### *TIMM50* disease-causing mutation reduces the levels of TIM23 complex components in patient fibroblasts

A previous study reported the clinical features of a rare genetic disease in several patients from two unrelated Arab families residing in Israel and Palestine (23). Patients suffering from the disease exhibited various neurological and metabolic disorders, including epilepsy, developmental delay, optic atrophy, cardiomyopathy, and 3-methylglutaconic aciduria. The disease was linked to a homozygous missense mutation resulting in T149M replacement in the *TIMM50* gene (Fig 1A). To unravel the molecular basis of the disease, we examined the effect of the mutation on the proteome of primary fibroblasts collected from two affected family members (P1 and P2), as well as from a healthy relative (HC).

**Fig 1.**
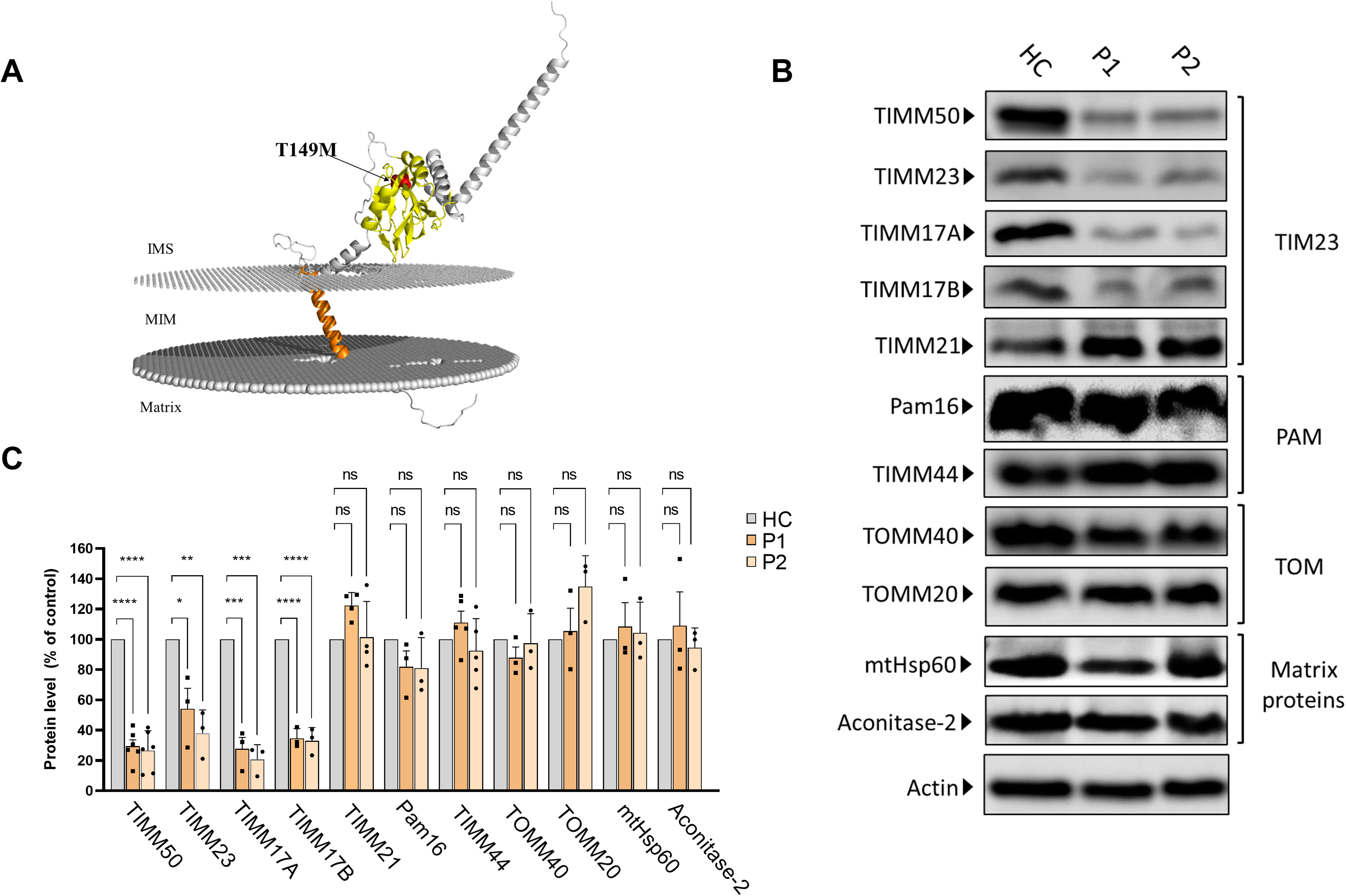
Differential effects of *TIMM50* mutation on the expression of TIM23, PAM and TOM subunits and matrix-destined proteins. (A) Predicted AlphaFold (57) human TIMM50 structure displaying the position of the T149M mutation. PPM webserver (58) was used to embed the predicted structure into the MIM. Orange – transmembrane domain, Yellow – FCP1-like domain, Red – Threonine 149. (B) Healthy control- and patients-derived primary fibroblasts were lysed and analyzed by immunoblot with the indicated antibodies. Actin was used as loading control. Full results and original blots are found in supplemental material S1 Raw images. (C) Band density analysis of the blots presented in A (and in supplemental material S1 Raw images), showing significant decrease in levels of TIMM50, TIMM23 and TIMM17A/B, but not in TIMM21, PAM subunits, TOM subunits and matrix TIM23 complex substrates. Band signals were normalized to the loading control and compared to the level measured in the healthy control sample, taken as 100%. Data are shown as means ± SEM, n = 6 biological repeats for TIMM50 antibody, n = 3-5 biological repeats for all other antibodies, *p-value < 0.05, **p-value < 0.01, ***p-value < 0.001, ****p-value < 0.0001, Ordinary one-way ANOVA.

We first examined the effect of the mutation on TIMM50 levels by immunoblotting and found that the mutation leads to a significant decrease in TIMM50 levels in patient fibroblasts (Fig 1B and C). We next examined the effect of the *TIMM50* mutation on other components of the TIM23 complex. Notably, our analysis revealed a significant reduction in the levels of the core TIM23 complex subunits, namely, TIMM23, TIMM17A and TIMM17B (Fig 1B and C). These results are in agreement with previous reports showing that two other *TIMM50* mutations (resulting in R217Q+G372S and S112*+G190A replacements, both compound heterozygous) also led to major decreases in TIMM50, TIMM23 and TIMM17A/B levels (24,25).

In contrast to the decreased levels of TIM23 complex core components seen in the patient fibroblasts, the levels of subunits belonging to the PAM complex were not affected, despite their expected import dependency on the TIM23 complex. TIMM21 also showed minimal to no change in amount. Additionally, as subunits of the TOM complex that serves as the general import pore in the outer membrane were shown to interact with TIM23 complex subunits, including TIMM50 (28), we also considered the possible effect of reduced TIMM50 levels on the expression of TOM complex subunits. No changes in the levels of TOM complex subunits addressed in patient fibroblasts were noted (Fig 1B and C).

Finally, as TIM23 is known to be the sole gateway into the mitochondrial matrix (7), we examined the effect of *TIMM50* mutation on the steady state levels of the matrix proteins aconitase 2 and mtHsp60. Surprisingly, the steady state levels of both proteins were unchanged in patient fibroblasts (Fig 1A and B), despite the significant loss of TIMM50, TIMM23, and TIMM17A/B.

### Generating a neuronal model system to study the neurophysiological effects of TIMM50 deficiency

As *TIMM50* mutations lead to severe neurodevelopmental symptoms and to a significant reduction in steady state TIMM50 levels (Fig 1 and (24,25)) we wanted to study the effects of TIMM50 deficiency in neuronal cells by knocking down TIMM50 in mouse primary cortical neurons. For this purpose, we designed three shRNA sequences (termed Sh1, Sh2 and Sh3; vector schemes are presented in Fig 2A) and cloned them into a lentiviral vector that also allows for EGFP expression under control of the hSyn promotor (29). These vectors allow TIMM50 knockdown (KD) while specifically labeling neurons with EGFP, allowing for efficient visualization in single cell experiments. We compared TIMM50 expression relative to three controls, namely, an untreated control (i.e., cultures that were not transduced), a pLL3.7 control (i.e., cultures that were transduced but did not transcribe the shRNA sequence, yet expressed EGFP), and a shRNA system activation control (i.e., cultures that were transduced with a scrambled shRNA sequence). All the three targeting shRNA sequences had an impact on TIMM50 levels, in comparison to the controls. Sh2 had the most significant and consistent effect, reducing TIMM50 levels by ∼80% (Fig 2B and C). Therefore, Sh2 was chosen as the KD vector for all subsequent experiments on neurons.

**Fig 2.**
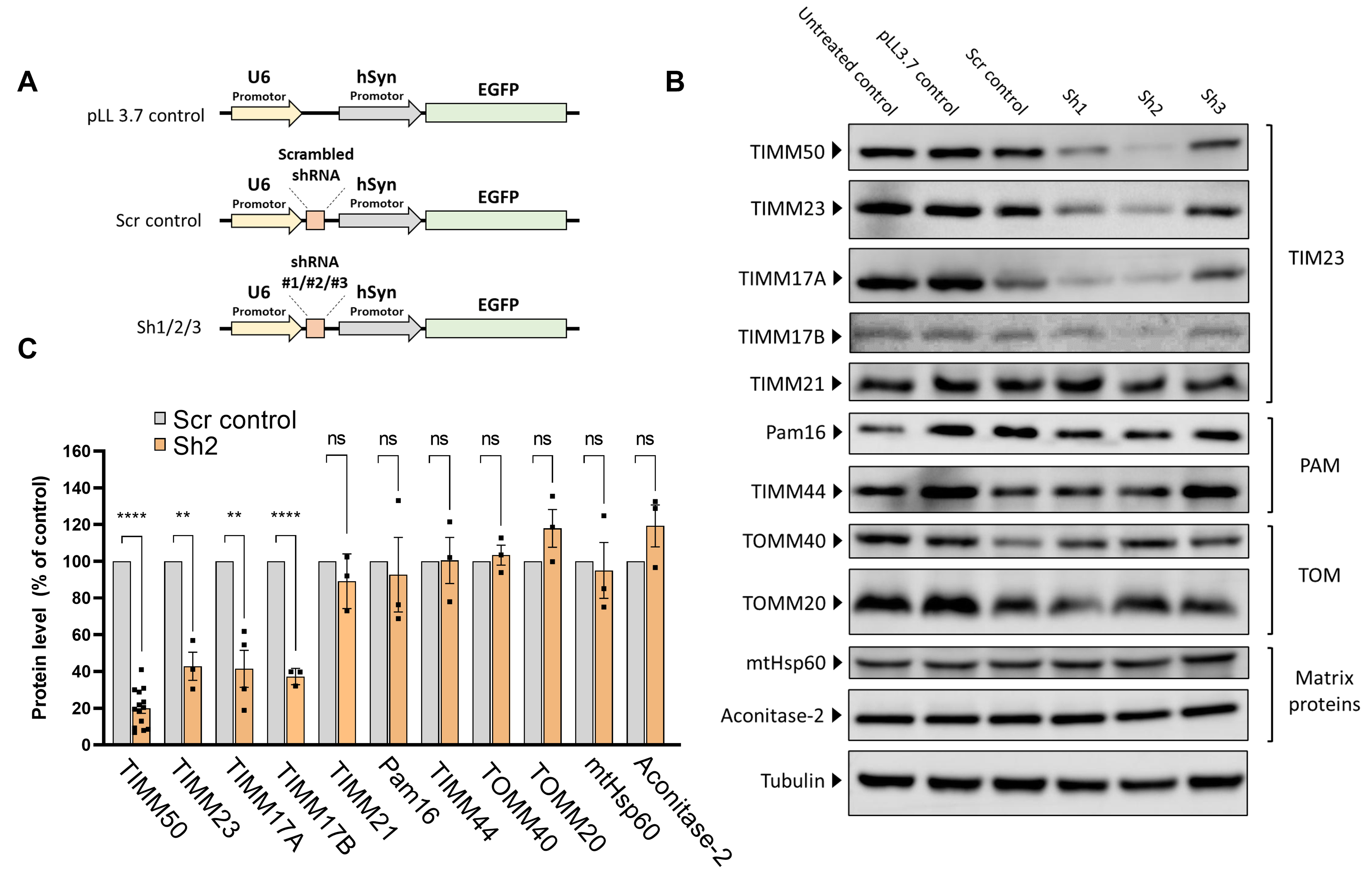
TIMM50 knockdown in mouse primary cortical neurons serves as a model system to study TIMM50 deficiency in mammalian neurons. (A) Schematic depictions of the different constructs used in this study. “pLL3.7 control” is a control for EGFP expression. “Scr control” is a control for the shRNA system activation with a scrambled Sh3 sequence and Sh1-Sh3 are three plasmids expressing different TIMM50 targeting shRNA sequences. (B) Neuronal cultures were transduced to express the indicated constructs, lysed and analyzed by immunoblot with the indicated antibodies. Tubulin was used as loading control. “Untreated” control are cells that were not transduced. Full results and original blots are found in supplemental material S2 Raw images. (C) Band density analysis of the blots presented in B (and in supplemental material S2 Raw images) showing significant decrease in levels of TIMM50, TIMM23 and TIMM17A/B, but not in TIMM21, PAM subunits, TOM subunits and matrix TIM23 complex substrates. Band signals were normalized to the loading control and compared to the Scr control, taken as 100%. Data are shown as means ± SEM, n = 14 biological repeats for TIMM50 antibody, n = 3-4 biological repeats for all other antibodies, **p-value < 0.01, ****p-value < 0.0001, unpaired Student’s t-test.

We next examined the effects of TIMM50 KD in neurons, addressing the same components as were tested in fibroblasts. In complete agreement with the fibroblast results, the levels of TIMM23 and TIMM17A/B were significantly decreased, while the amounts of TIMM21, PAM subunits, TOM subunits and the matrix substrates tested were not affected (Fig 2B and C). Hence, similarly to what was detected in fibroblasts, steady state levels of several representative mitochondrial proteins were not affected in TIMM50 KD neurons.

### TIMM50 deficiency does not affect the steady state levels of a majority of its substrates

It was expected that the significant decrease in the levels of TIM23 core components seen upon TIMM50 deficiency would decrease the levels of substrates processed by this translocation complex. The observation that steady state levels of two mitochondrial matrix substrates, mtHsp60 and aconitase 2, were not affected by the significant loss of TIM23 core components in both research systems (Fig 1A and B and Fig 2B and C), motivated us to examine the impact of TIMM50 KD on the total mitochondrial and cellular proteomes. For this purpose, we performed untargeted mass spectrometry analysis of fibroblasts from both patients, as compared to the healthy control, and of mice primary neurons transduced with either Sh2 or the Scr control. For fibroblasts, principal component analysis (PCA) was performed to verify that the replicates of both patients are consistently different from the HC replicates (Fig S1).

In the case of fibroblasts, 127 MIM and 190 matrix proteins were detected using mass spectrometry (Supplemental material S1 and S2 Datasets). Surprisingly, we noticed that the levels of 83 (∼65%) MIM proteins and 135 (∼71%) matrix proteins were not affected in either patient, as compared to the HC (Fig 3A and B and supplemental material S1 and S2 Datasets). Among the MIM proteins that were not affected by the *TIMM50* mutation, we identified multiple proteins involved in calcium homeostasis (such as MICU2, SLC25A3 and LETM1), heme synthesis (such as PPOX and CPOX), and cardiolipin synthesis (HADHA). Among matrix proteins that were not affected by *TIMM50* mutation, we identified multiple proteins involved in Fe-S cluster biosynthesis (such as NFS1, GLRX5 and ISCU), detoxification (such as PRDX5, SOD2, ABHD10 and GSTK1), fatty acid oxidation (such as DECR1, ECHS1 and ETFA), and amino acid metabolism (PYCR1, ALDH18A1 and HIBCH). Also, the majority of TCA cycle proteins, such as ACO2, DLST, IDH3B and OGDH, found in the matrix, were not affected in patient fibroblasts. Interestingly, every MIM protein that changed significantly in both patients, appeared to be downregulated, while the significantly changed matrix proteins comprised of both downregulated and upregulated proteins (Fig 3C). Unexpectedly, a few matrix proteins (ALDH2, GRSF-1, AK4, LACTB2 and OAT) showed increased steady state levels in both patients (Fig 3C). Of these proteins, ALDH2 and GRSF-1 showed ∼15 and ∼6-fold increases, respectively, as was also confirmed by immunoblot (Fig S2A).

**Fig 3.**
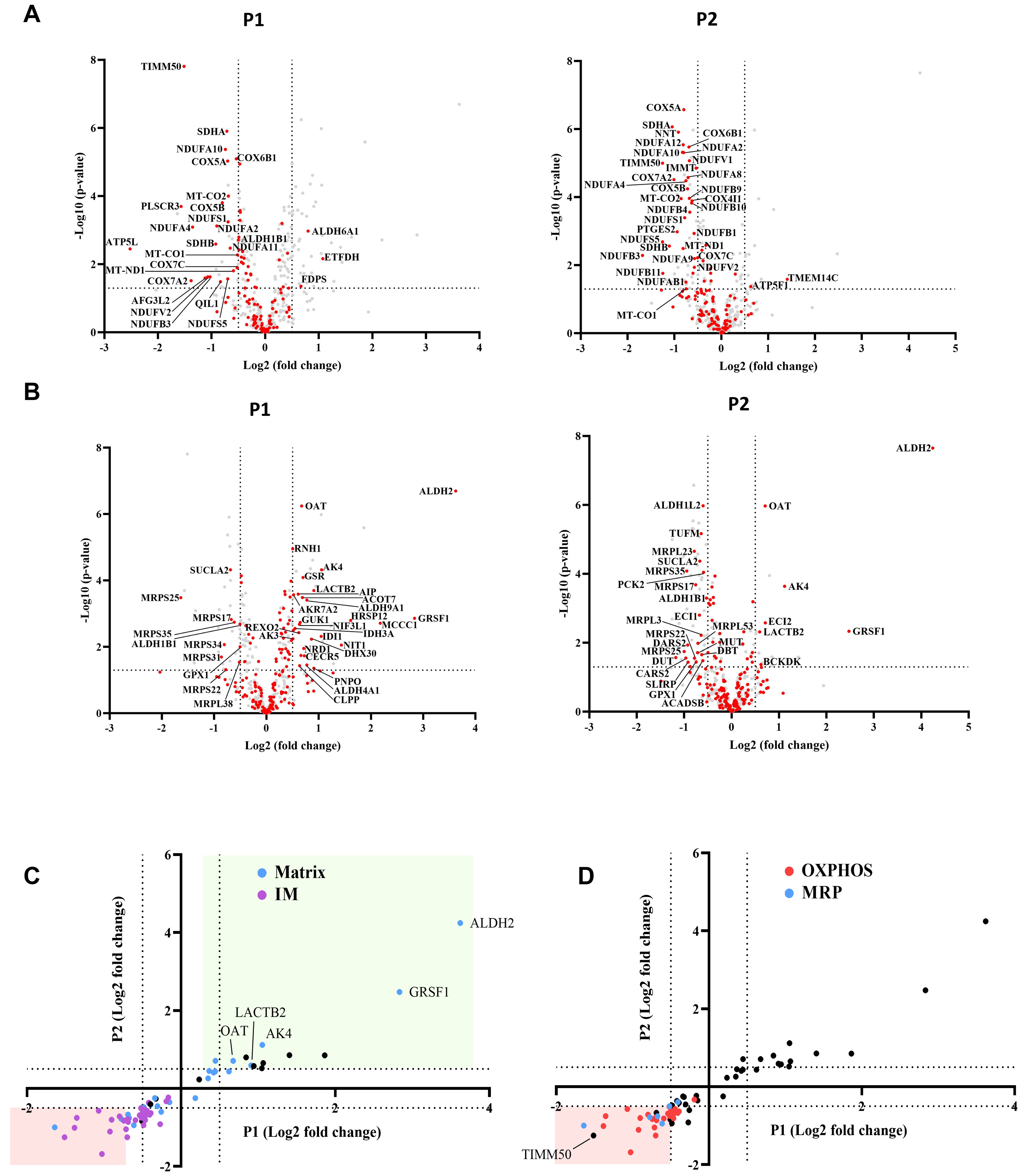
TIMM50 deficiency leads to a specific reduction in the OXPHOS and MRP machineries but does does not affect the majority of MIM and mitochondrial matrix proteome. Untargeted mass spectrometry analysis of fibroblasts from both patients (P1, P2), as compared to the healthy control. In the volcano plots displayed in (A, B) and in all subsequent figures, the y-axis cut off of >1.301 corresponds to –log (0.05) or p-value = 0.05, while the x-axis cut off of <−0.5 and >0.5 corresponds to a ±1.414-fold change. Each dot in the graph represents a protein. Proteins depicted on the right side of the x-axis cut-off and above the y-axis cut off were considered to be increased in amount, while proteins depicted on the left side of the x-axis cut-off and above the y-axis cut off were considered to be decreased in amount. All relevant proteins that were increased/decreased in amount are identified by their gene name, next to the dot. Statistical analysis was performed using Student’s t-test and a p-value <0.05 was considered statistically significant. n = 9 per group (three biological repeats in triplicate), P1 and P2 results were compared to HC results. Full list of differentially expressed proteins in fibroblasts is found in supplemental material S1-2 Datasets. (A) The steady state levels of a majority of MIM proteins detected in patient fibroblasts were not affected. MIM proteins are colored red, while other detected mitochondrial proteins are colored grey. (B) The steady state levels of a majority of matrix proteins detected in patient fibroblasts were not affected. Matrix proteins are colored red, while other detected mitochondrial proteins are colored grey. (C) Correlation plot comparing the log-fold-change of P1 to that of P2 showing that out of the proteins that were significantly changed for both patients, MIM proteins are consistently downregulated, while matrix proteins are both upregulated and downregulated. (D) Correlation plot comparing the log-fold-change of P1 to that of P2 showing that OXPHOS and MRP proteins comprise nearly all the proteins that were downregulated in both patients.

### Oxidative phosphorylation subunits and mitochondrial ribosomal proteins comprise the majority of down-regulated proteins in TIMM50-deficient fibroblasts

A total of 69 oxidative phosphorylation (OXPHOS) and 27 mitochondrial ribosomal proteins (MRP) were detected in fibroblasts by mass spectrometry. Remarkably, of the 18 MIM proteins found in decreased amounts in both patients, 17 belong to the OXPHOS family, while of the 7 matrix proteins found in decreased amounts in both patients, 4 proteins belong to the MRP family (Fig 3D). The OXPHOS complexes most affected were CI, CII and CIV. In the case of CI, one membrane core subunit (MT-ND1), two hydrophilic core subunits (NDUFS1 and NDUFV2), and four super-numerary subunits (NDUFA2, NDUFA10, NDUFB3, and NDUFS5) exhibited a significant negative fold-change in both patients (supplemental material S1 and S2 Datasets). The levels of two CII subunits, namely, SDHA and SDHB, and eight CIV subunits, including the two catalytic subunits MT-CO1 and MT-CO2, were also significantly decreased in both patients. GO term analysis performed on all the genes that were downregulated in both patients, confirmed enrichments of mitochondrial inner membrane proteins, several OXPHOS related processes and mitochondrial ribosome subunits (Fig S3A).

Interestingly, as mentioned above, several mitochondrially encoded OXPHOS subunits were found to be downregulated. These subunits are translated by the mitochondrial ribosomes and assembled into the MIM by the oxidase assembly (OXA) insertase (30). Therefore, these subunits are not directly related to TIM23 import. To elucidate the reason behind downregulation of these proteins, we tested the levels of the OXA insertase by immunoblotting and found no significant difference in its levels between the patient fibroblasts and the HC (Fig S3B and C). Therefore, we conclude that the decreased levels of MRP subunits likely lead to lower translation rates of mitochondrially encoded OXPHOS subunits and consequently, to their lower abundance in the MIM.

### The impact of TIMM50 deficiency on the mitochondrial proteome in neurons

In TIMM50 KD neurons, 170 MIM and 215 matrix proteins were detected by mass spectrometry (supplemental material S3 Dataset). Similar to the observations for fibroblasts, majority of the MIM and matrix proteins were not affected by TIMM50 KD (Fig. 4A and B). OXPHOS and MRP subunits showed a similar trend to what was observed in patient fibroblasts and appeared to be mostly downregulated, although the affected OXPHOS subunits appeared to decrease to a lesser extent (Fig 4C and D). The difference in the extent of fold-change in neurons, as compared to patient fibroblasts, could be due to the short duration of KD in the neuronal experiment, as compared to that with the patient fibroblasts, that constantly carry the deficiency.

**Fig 4.**
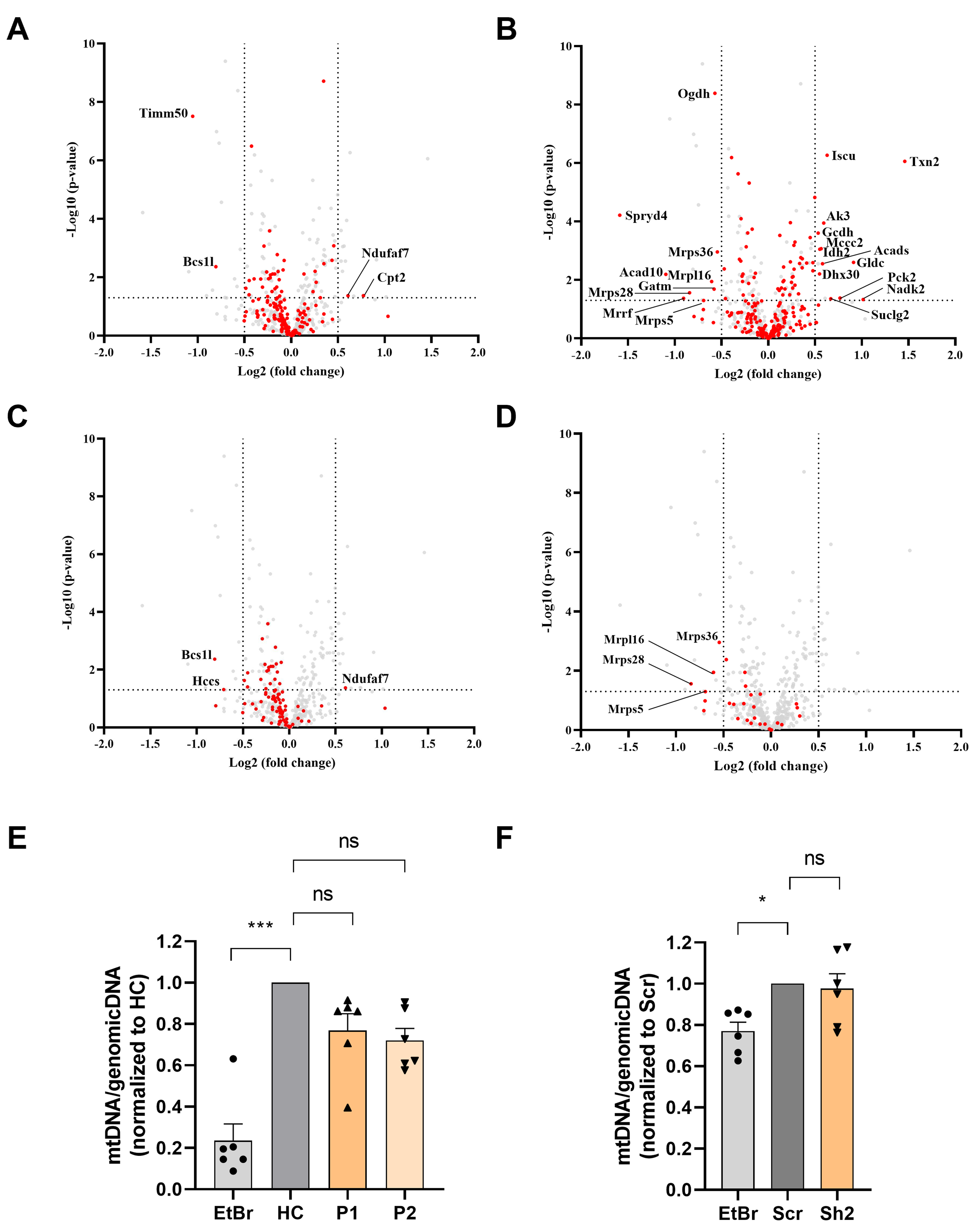
The impact of TIMM50 deficiency on the mitochondrial proteome in neurons. Untargeted mass spectrometry analysis of mice primary neurons transduced with either Sh2 or the Scr control. n = 9 per group (three biological repeats in triplicates). Full list of differentially expressed proteins in neurons is found in supplemental material S3 Dataset. (A) The steady state levels of a majority of MIM proteins detected in TIMM50 KD neuronal cells were not affected. MIM proteins are colored red, while other detected mitochondrial proteins are colored grey. (B) The steady state levels of a majority of matrix proteins detected in TIMM50 KD neuronal cells were not affected. Matrix proteins are colored red, while other detected mitochondrial proteins are colored grey. (C) Most OXPHOS subunits are observed to be in decreased amounts in TIMM50 KD neuronal cells, albeit at a lower extend from what was observed in patient fibroblasts. OXPHOS proteins are colored red, while other detected mitochondrial proteins are colored grey. (D) Decreased steady state levels of multiple MRP subunits was observed in TIMM50 KD neuronal cells. MRP proteins are colored red, while other detected mitochondrial proteins are colored grey. (E, F) Mitochondrial DNA content in fibroblasts (E) and neurons (F) was not affected in TIMM50 deficient cells compared to control cells. Mitochondrial DNA content was estimated by measuring the ratio between mitochondrial and nuclear DNA. Dloop1 expression was measured by qPCR relative to HPRT; TERT served as a control of a nuclear-encoded gene. Ethidium bromide (EtBr; 100 ng/ml) was used as positive control. Data are shown as means ± SEM, n = 6 (samples from three biological replicates, each performed twice), *p-value < 0.05, ***p-value < 0.001, Kruskal-Wallis test.

Additionally, to determine whether the changes observed in the levels of OXPHOS and MRP proteins were an indirect effect resulting from general mitochondrial DNA loss, we performed qPCR assessment of the mitochondrial DNA content. Our results showed no effect on mitochondrial DNA levels in either model system (Fig 4E and F), thereby confirming that the observed effects on the OXPHOS and MRP protein levels were the direct and specific result of TIMM50 deficiency, and not indirectly due to mitochondrial DNA loss.

Overall, we conclude that translocation of the majority of MIM and matrix proteins was not affected in both patient fibroblasts and TIMM50 KD mice neurons, even after significant disruption of TIMM50 and the TIM23 core subunits. However, TIMM50 deficiency severely affected two major mitochondria complex systems, namely, the OXPHOS and MRP protein machineries.

### TIMM50 deficiency affects ATP production

The observed decrease in steady state levels of OXPHOS subunits led us to examine oxygen consumption in TIMM50-deficient cells. For this purpose, we performed the Seahorse XF cell Mito Stress test (Fig 5A). Comparing oxygen consumption rates in the HC, P1 and P2 fibroblasts, and in the Sh2- and Scr control-transduced neuronal cells revealed significant impairment of both basal and maximal respiration rates and of OXPHOS-dependent energy production rate in the non-control cells (Fig 5C and D). Overall, these results suggest that TIMM50 deficiency severely affects the OXPHOS and MRP protein machineries and leads to OXPHOS-dependent ATP deficiency in both systems.

**Fig 5.**
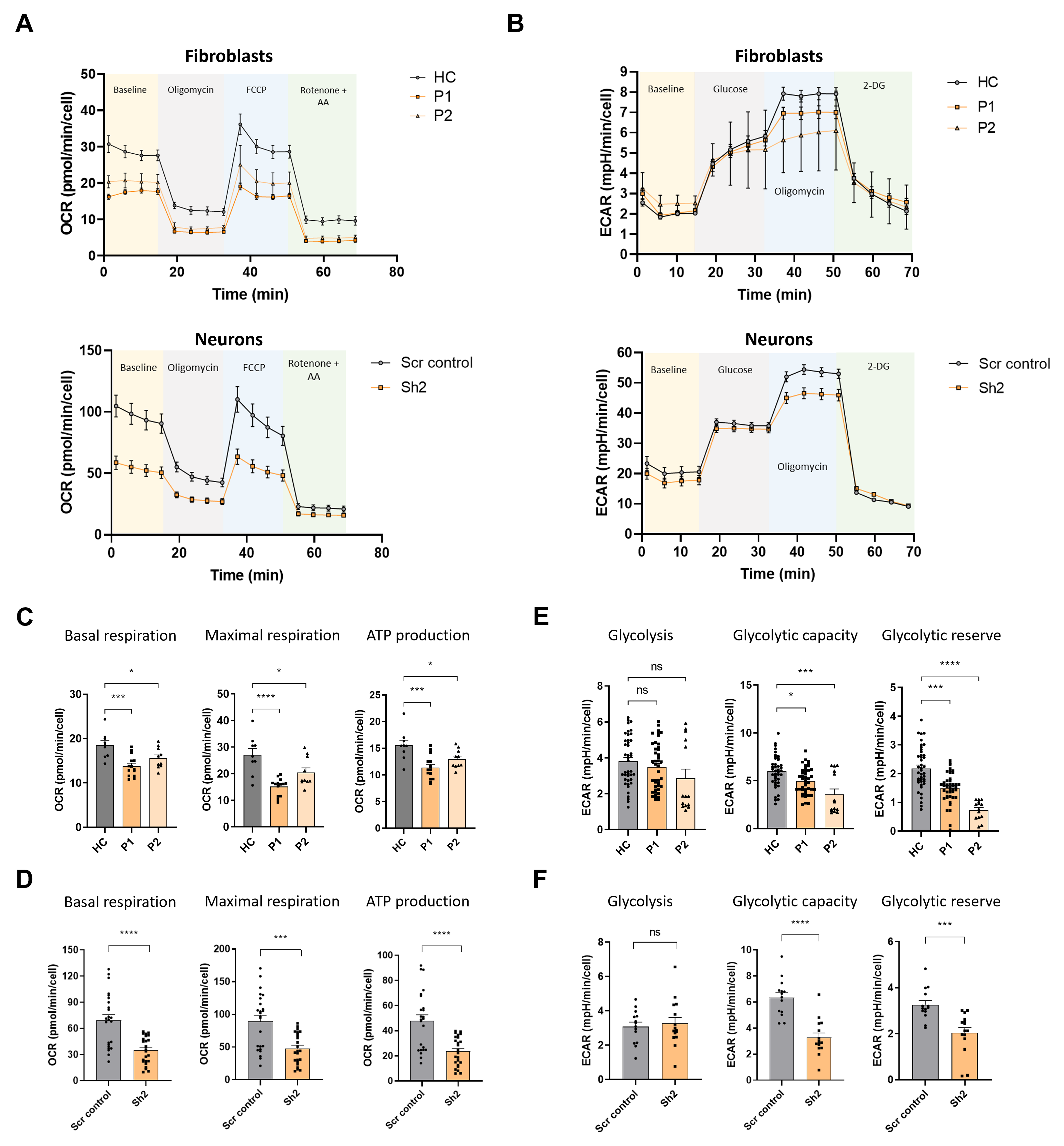
TIMM50 deficiency negatively impacts OXPHOS machinery and glycolysis functions. (A) A Seahorse XF cell mito-stress assay was used to measure mitochondrial oxygen consumption rates at basal levels and in response to the indicated effectors in fibroblasts (upper panel) and neuronal cells (lower panel). (B) A Seahorse XF cell glycolysis stress assay was used to measure glycolysis at basal levels and in response to the indicated effectors in fibroblasts (upper panel) and neuronal cells (lower panel). (C) Basal respiration, maximal respiration and ATP-linked respiration were reduced in patient fibroblast cells compared to HC cells. Data are shown as means ± SEM, *p-value < 0.05, ***p-value < 0.001, ****p-value < 0.0001, Ordinary one-way ANOVA. (D) Basal respiration, maximal respiration and ATP-linked respiration were reduced in TIMM50 KD neuronal cells compared to Scr control-transduced neuronal cells. Data are shown as means ± SEM, ***p-value < 0.001, ****p-value < 0.0001, unpaired Student’s t-test. (E) Basal glycolysis remained similar, while glycolytic capacity and glycolytic reserves were reduced in patient fibroblast cells compared to HC cells. Data are shown as means ± SEM, *p-value < 0.05, ***p-value < 0.001, ****p-value < 0.0001, Kruskal-Wallis test. (F) Basal glycolysis remained similar, while glycolytic capacity and glycolytic reserves were reduced in TIMM50 KD neuronal cells compared to Scr control-transduced neuronal cells. Data are shown as means ± SEM, ***p-value < 0.001, ****p-value < 0.0001, unpaired Student’s t-test. For all the experiments shown in (A-F), n = 4-6 biological repeats of 3-6 technical repeats each.

Moreover, as cells are able to meet their energetic requirements via glycolysis when the OXPHOS apparatus malfunctions (31), we measured the glycolytic capabilities of TIMM50-deficient cells by performing a Seahorse XF glycolysis stress test (Fig 5B). In both systems, the basal glycolysis level remained stable, in comparison to controls, suggesting that the cells did not make the metabolic switch so as to increasingly rely on the non-mitochondrial energy production pathway that is glycolysis (Fig 5E and F, left panel). Moreover, both glycolytic capacity and glycolytic reserves were significantly reduced, indicative of an impaired ability of both patient fibroblasts and neurons to switch their energetic emphasis to glycolysis when needed (Fig 5E and F, middle and right panels).

### TIMM50 KD impairs mitochondrial trafficking in neuronal cells

The transport of mitochondria within neuronal processes is crucial for cell survival (32–34). Therefore, we investigated the effect of TIMM50 deficiency on mitochondrial trafficking in neuronal cells. To track individual mitochondria in neuronal processes, we used a transfection method, instead of transduction, that results in low expression efficiency in neurons (35). This allowed us to visualize individual processes and track single mitochondria with minimal background noise. To visualize neuronal cells and their mitochondria, we co-transfected our cultures with a dsRed-mito-encoding plasmid, together with the KD or control plasmids. We then performed live cell imaging of individual neuronal processes and tracked the movement of individual mitochondria in these structures (for examples, see Fig 6A and S1 Movie). The live imaging sets were converted into kymographs and calibrated in time and space, which allowed extraction of different trafficking parameters, such as distance of movement, speed, and percentage of moving mitochondria (Fig 6B).

**Fig 6.**
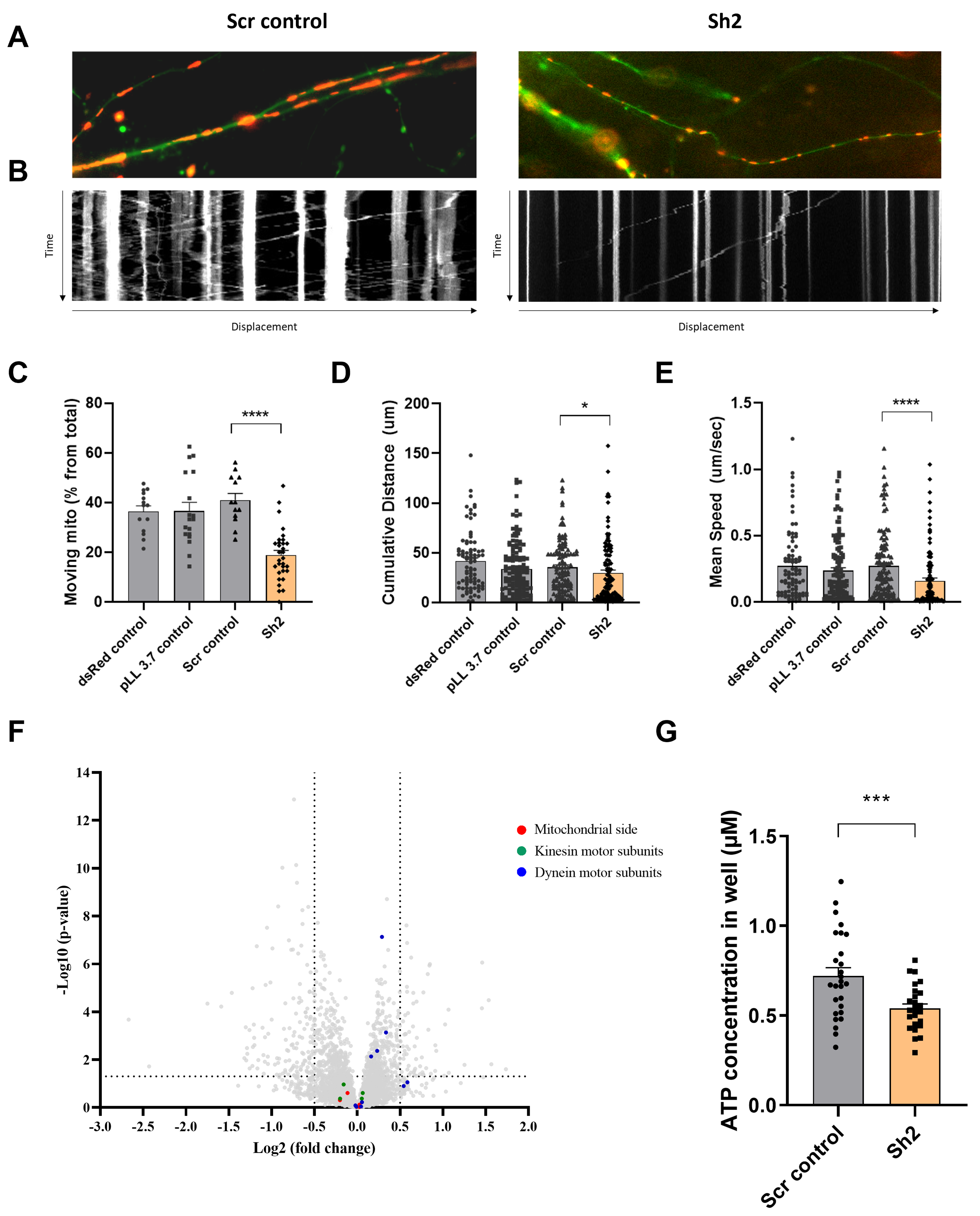
TIMM50 deficiency leads to defective mitochondrial trafficking along neuronal processes. (A) Representative images of neuronal processes in neuronal cultures that were co-transfected with a dsRed-mito plasmid and either a Scr control plasmid (left panel) or a TIMM50 KD plasmid (right panel). (B) Kymographs of the same processes in A, showing the displacement of mitochondria over time. Y axis length is 5 minutes, X axis length is about 100 μm. (C) Lower percentage of moving mitochondria was observed in TIMM50 KD neuronal processes compared to controls. Each dot in the graph represents the percentage of moving mitochondria out of the total observed mitochondria in a single neurite. Data are shown as means ± SEM, n = 13-31 neurites per condition, analyzed from three biological repeats, ****p-value < 0.0001, Ordinary one-way ANOVA. (D) Decreased mitochondrial cumulative travelling distance was observed in TIMM50 KD neuronal cells compared to controls. Each dot in the graph represents a single moving mitochondrion. n = 84-130 mitochondria per condition, analyzed from three biological repeats, *p-value < 0.05, Kruskal-Wallis test. (E) Slower mitochondrial movement was observed in TIMM50 KD neuronal cells compared to controls. Each dot in the graph represents a single moving mitochondrion. n = 84-130 mitochondria per condition, analyzed from three biological repeats, ****p-value < 0.0001, Kruskal-Wallis test. (F) No change in mitochondrial trafficking proteins was observed TIMM50 KD neuronal cells. The y-axis cut off of >1.301 corresponds to –log (0.05) or p-value = 0.05, while the x-axis cut off of <−0.5 and >0.5 corresponds to a ±1.414-fold change. Each dot in the graph represents a protein. Proteins depicted on the right side of the x-axis cut-off and above the y-axis cut off were considered to be increased in amount, while proteins depicted on the left side of the x-axis cut-off and above the y-axis cut off were considered to be decreased in amount. Statistical analysis was performed using Student’s t-test and a p-value <0.05 was considered statistically significant. n = 9 per group (three biological repeats in triplicate). Full list of differentially expressed proteins in neurons is found in supplemental material S3 Dataset. (G) Lower cellular ATP levels were observed in TIMM50 KD neuronal cells compared to Scr control-transduced cells. Data are shown as mean ± SEM, n = 27 quantified wells (each containing 5×10^4^ cells) per condition, from three biological repeats with nine technical repeats each, ***p-value < 0.001, unpaired Student’s t-test.

Our results showed a two-fold decrease in the percentage of mobile mitochondria in TIMM50-deficient neuronal cells, as compared to control cells (Fig 6C). Moreover, mobile mitochondria in TIMM50-deficient neuronal cells tend to cover less distance and travel at a lower average travelling speed (Fig 6D and E). This indicates that TIMM50 deficiency causes neuronal cell mitochondria to be more static, which could consequently lead to further energy deprivation in regions where mitochondria are needed but cannot be shipped.

Mitochondrial movement along neuronal processes is coordinated by a motor/adaptor complex. The motors kinesin and dynein use ATP to move organelles along microtubules. Mitochondria are assembled onto these motors via a mitochondrial outer membrane protein called Miro, a cytosolic adaptor called Milton (also known as TRAK1/2), and a few cytosolic accessory proteins (36). Although TIM23 and TIMM50 are not directly involved in the biogenesis of any of these proteins, we, nonetheless, examined their expression levels following TIMM50 KD. Our proteomics data revealed no major changes in the levels of proteins involved in mitochondrial trafficking (Fig 6F), suggesting that the observed effect on neuronal cell mitochondrial trafficking to be most likely indirect and resulting from the ATP deficiency. To examine this possibility, we measured cellular ATP levels in Scr control-transduced and TIMM50 KD neuronal cells. As expected from the impaired ATP production (Fig. 5), we found that TIMM50 KD led to a significant reduction of about 25% in cellular ATP levels (Fig 6G).

### TIMM50 KD leads to excess neuronal activity and increased action potential frequency

All TIMM50 mutant patients studied thus far displayed severe neurological pathologies, which include epilepsy, developmental delay, and loss of movement abilities. Such abnormalities can be attributed to alternations in basic neuronal function. To assess whether such functions are altered upon TIMM50 KD in neuronal cells, we measured intrinsic neuronal excitability, as well as spontaneous neurotransmitter release, using the whole cell patch clamp technique.

Initially, we measured spontaneous excitatory activity of the cells in the presence of tetrodotoxin (TTX) (representative traces are presented in Fig S4A). We quantified the average miniature excitatory post-synaptic current (mEPSC) amplitude, area and frequency in each of the measured neuronal cells and found no significant differences between any of these measures in KD cells, as compared to controls (Fig S4B). Moreover, the relative frequency distribution of the amplitude measurements showed an even distribution pattern (Fig S4C). Examination of the cumulative distribution function confirmed that no significant differences in mEPSC amplitude distribution exists in the neuronal cells (Fig S4D).

We subsequently measured the minimal current required to induce an action potential by slowly increasing the stimulation current in a stepwise manner (Fig 7A). This allowed us to estimate the rheobase of the neuronal TIMM50 KD and control cells. We found that there were no significant differences in the current needed to trigger an action potential between the TIMM50 KD and Scr control-transduced cells (Fig 7D). We also assessed characteristics of the first observed action potential in each measurement. Both the half-width and rate of fall of the first action potential were similar in TIMM50 KD and Scr control-transduced neurons (Fig 7B, E and F). However, TIMM50 KD cells displayed shorter action potential latency (Fig 7G) and a significant increase in the maximum number of action potentials as compared to the Scr control-transduced cells (Fig 7C and H). Overall, these results suggest that TIMM50 KD causes the cells to fire more action potentials without decreasing the firing threshold, probably due to a faster recovery time between successive action potentials.

**Fig 7.**
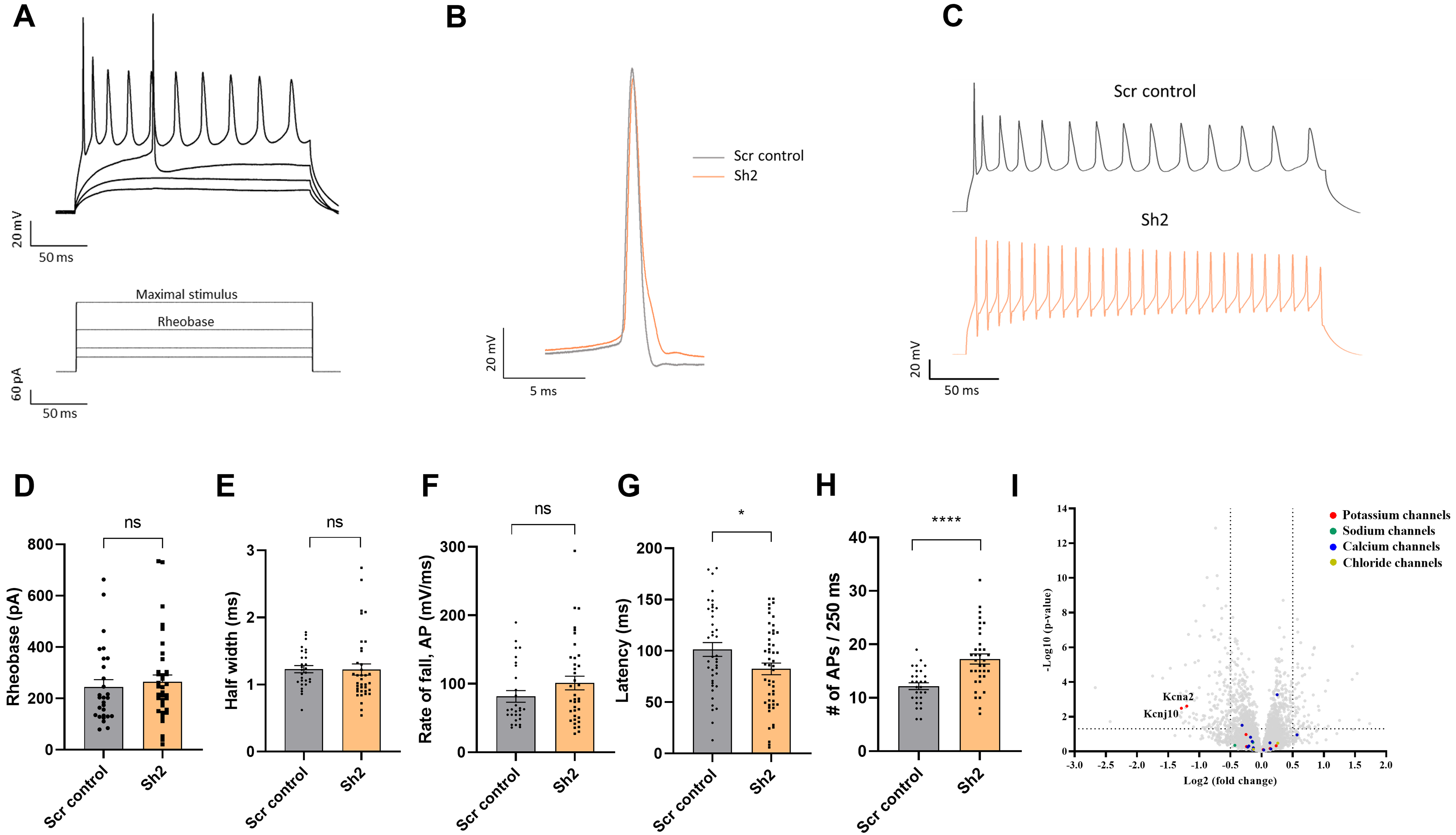
TIMM50 deficiency leads to a significant decrease in the levels of KCNA2 and KCNJ10 potassium channels and an increased electrical activity. (A) Example of voltage traces in response to increased depolarization of TIMM50 KD neurons. The bottom panel shows the protocol applied, while the top panel shows the typical response pattern that was measured. (B) Representative traces of a single action potential, received at rheobase level, from a TIMM50 KD neuron, as compared to Scr control-transduced neuron. (C) Representative traces of the maximal stimulus for each condition, showing the maximal amount of action potentials measured for each group. (D) No change in rheobase was observed in TIMM50 KD neuronal cells, as compared to Scr control-transduced cells. Rheobase was measured as the first current step that caused firing of an action potential. Data are shown as means ± SEM, n = 28 / 37 cells (for Scr control-/-Sh2-transduced cells, respectively) from three biological repeats, Mann-Whitney test. (E) No change in the action potential half-width was observed in TIMM50 KD neuronal cells, as compared to Scr control-transduced cells. Half-width was measured as the time between the rising and falling phases of the action potential, at the half-point between the tip of the peak and the bottom of the voltage rising curve. Data are shown as means ± SEM, n = 28 / 37 cells (for Scr control-/ Sh2-transduced cells, respectively) from three biological repeats, Mann-Whitney test. (F) No change in the action potential rate of fall was observed in TIMM50 KD neuronal cells, as compared to Scr control-transduced cells. Rate of fall was measured as the time between the action potential peak and the baseline following the action potential, divided by the change in voltage between the same two points (ΔX/ΔY). Data are shown as means ± SEM, n = 28 / 37 cells (for Scr control / Sh2 respectively) from three biological repeats, Mann-Whitney test. (G) Decreased action potential latency was observed in TIMM50 KD neuronal cells compared to Scr control-transduced cells. Latency was measured as the time difference between the beginning of the pulse to the peak of the action potential. Each dot in the graph represents the average latency of the first five consecutive action potentials appearing after rheobase level. Data are shown as means ± SEM, n = 40 / 50 cells (for Scr control-/Sh2-transduced cells, respectively) from four biological repeats, *p-value < 0.05, unpaired Student’s t-test. (H) An increase in the maximal number of action potentials fired in a single stimulus was observed in TIMM50 KD neuronal cells, as compared to Scr control-transduced cells. Data are shown as means ± SEM, n = 28 / 37 cells (for Scr control-/Sh2-transduced cells, respectively) from three biological repeats, ****p-value < 0.0001, unpaired Student’s t-test. (I) Amongst the detected ion channel proteins, a specific decrease in KCNA2 and KCNJ10 potassium channels was observed in TIMM50 KD neuronal cells. The y-axis cut off of >1.301 corresponds to –log (0.05) or p-value = 0.05, while the x-axis cut off of <−0.5 and >0.5 corresponds to a ±1.414-fold change. Each dot in the graph represents a protein. Proteins depicted on the right side of the x-axis cut-off and above the y-axis cut off were considered to be increased in amount, while proteins depicted on the left side of the x-axis cut-off and above the y-axis cut off were considered to be decreased in amount. Statistical analysis was performed using Student’s t-test and a p-value <0.05 was considered statistically significant. n = 9 per group (three biological repeats in triplicate). Full list of differentially expressed proteins in neurons is found in supplemental material S3 Dataset.

An increase in the firing frequencies of action potentials can be explained by a reduced presence of voltage-dependent potassium channels, which are known to control spike frequency (37). Indeed, our proteomics results confirmed a decrease of about 2.5-fold in the levels of the KCNA2 and KCNJ10 potassium channels in the TIMM50-deficient neuronal cells, supporting the increase in their action potential frequency (Fig 7I). KCNA2 levels were also tested via immunoblot and confirmed to be dramatically decreased (Fig S2B). To further test how KCNA2 reduction impacts cellular electrical activity, we used α-dendrotoxin (α-DTX), a known KCNA2 channel blocker (38,39), to mimic a reduction in KCNA2. The number of action potentials fired was measured before and after application of 100 nM α-DTX to Scr control-transduced and TIMM50 KD neuronal cells (Fig S4E). As expected, the difference in the number of action potentials after and before α-DTX treatment increased significantly more in Scr control-transduced cells than in TIMM50 KD cells (Fig S4F), given how TIMM50 KD cells initially contain less KCNA2 channels than do the corresponding controls. These data indicate that a reduction in KCNA2 contributes to the observed increase in firing rate in TIMM50 KD neurons.

## Discussion

The TIMM50 protein is a pivotal member of the TIM23 complex that is suggested to participate in the import of nearly 60% of the mitochondrial proteome (5,6). Human *TIMM50* mutations lead to neurological effects, including mitochondrial epileptic encephalopathy, intellectual disability, seizure disorders like infantile spasms, and severe hypotonia accompanied by 3-methylglutaconic aciduria (22–27). Despite being involved in the import of the majority of the mitochondrial proteome, little is known about the effects of TIMM50 deficiency on the entire mitochondrial proteome. Additionally, despite the fact that human TIMM50 mutations lead mainly to neurological symptoms, no study has yet addressed the effects of TIMM50 deficiency in brain cells. Therefore, in this study, we utilized two research models – TIMM50 mutant patient fibroblasts and TIMM50 KD primary mouse neuronal cultures, to study the impact of TIMM50 deficiency on the mitochondrial proteome and its impact on neurophysiology.

Interestingly, in both model systems, TIMM50 deficiency reduced the levels of TIM23 core subunits, yet did not alter steady state levels of a majority of TIM23 substrates. These observations are similar to the recent analysis of patient-derived fibroblasts which demonstrated that *TIMM50* mutations lead to severe deficiency in the level of TIMM50 protein (25,40). Notably, this decrease in TIMM50 was accompanied with a decrease in the level of other two core subunits, TIMM23 and TIMM17. However, unexpectedly, proteomics analysis in our study and that conducted by Crameri et al., 2024 indicate that steady state levels of most TIM23-dependent proteins are not affected despite a drastic decrease in the levels of the TIM23^CORE^ complex (40). The most affected proteins constitute of intricate complexes, such as OXPHOS and MRP machineries. Thus, both these studies might indicate that even reduced levels of the TIM23^CORE^ components are sufficient for maintaining the steady state levels of most presequence containing substrates. This is surprising, as normal TIM23 complex levels are suggested to be indispensable for the translocation of presequence-containing mitochondrial proteins (14,15,41–43). Even more surprising was that the amounts of some TIM23 substrates related to intricate metabolic and maintenance activities (e.g., ALDH2, GRSF-1, OAT, etc.) were increased.

These observations can be explained by several plausible mechanisms: it is possible that unlike what occurs in yeast, normal levels of mammalian TIMM50 and TIM23 complex are mainly essential for maintaining the steady state levels of intricate complexes/assemblies. Another explanation for this scenario is that the normal quantities of unaffected matrix proteins might be low, and hence, even ∼10-20% of functional TIMM50 protein might be sufficient to maintain their steady state levels. Alternatively, the presequence of such proteins might contain mitochondrial targeting signals (MTSs) that receive priority over other presequence-containing precursor proteins, thus enabling their translocation even in the presence of very few functional TIM23 complexes. However, further experiments examining these possibilities are needed for understanding the compromised TIM23-mediated protein import.

As stated earlier, loss of TIMM50 led to a significant reduction in the steady state levels of other TIM23 core subunits (i.e., TIMM23 and TIMM17A/B). Notably, TIMM23 and TIMM17A/B are thought to be imported by the TIM22 complex (44). However, we observed that the steady state levels of TIM22 complex subunits had not been affected by TIMM50 deficiency (supplemental material S1-3 Datasets). This indicates a hitherto unknown relation between steady state levels of TIMM50 and other TIM23 core subunits, which might affect the complex assembly process. Various structural and biochemical studies have attempted to elucidate the intricate structure of the TIM23 complex and the complicated interactions between its different subunits (11,12,45), however, little is presently known of the dynamic assembly processes of the complex. The fact that TIMM50 KD leads to a major reduction in the levels of TIMM23 and TIMM17A/B, despite a lack of direct dependency of the import of these proteins on TIMM50, and does not affect the levels of other TIM23 complex subunits, could provide a basis for future studies examining the assembly process of import complexes.

Our proteomics data and Seahorse XF analysis, paired with cellular ATP measurements, indicated that lower ATP levels were present in TIMM50-deficient cells. Specifically in the case of neurons, such ATP deficiency could be responsible for the negative impact seen on mitochondrial trafficking in neuronal cell processes, which could contribute to the various neurodegenerative phenotypes linked to the TIMM50 disease. Additionally, the detected increase in action potential firing rate in TIMM50-deficient neurons can explain the presence of epileptic seizures, a hallmark of all TIMM50 patients studied thus far. A plausible explanation for the increased action potential firing rate could be the 2.5-fold reduction in the levels of the KCNA2 (K_v_1.2) and KCNJ10 (K_ir_4.1) voltage dependent potassium channels, as revealed by our proteomics and immunoblot analysis. The KCNA2 (K_v_1.2) channel is a slowly inactivating channel that regulates neuronal excitability and firing rate (46). In peripheral sensory neurons, K_v_1.2 helps to determine spike frequency, while in neurons of the medial trapezoid body, the Kv1.1, Kv1.2, and Kv1.6 subunits are important regulators of repetitive spiking (39,47). While the inactivation of several of the potassium channels like K_v_1.2, K_v_1.1 and Kcnj10 (K_ir_4.1) are important for neuronal excitability, action potential width, and firing properties, in general, mutations in the genes encoding these channels that alter their inactivation are known to lead to temporal lobe epilepsy (46,48,49). In addition, blocking the activation of the K_v_1.2 by α-DTX had no effect on the resting membrane potential and only small effects on the amplitude and duration of the action potential (39), similar to what we observed in TIMM50 KD neurons, that showed a reduction in K_v_1.2 levels. Furthermore, blocking K_v_1.2 activation by α-DTX increased the frequency of action potentials in visceral sensory neurons (39), which, again, agrees with our observation that α-DTX application increased the firing rate in Scr control-transduced neurons, while hardly affecting the firing rate of TIMM50 KD neurons as they express lower levels of K_v_1.2. Hence, similar changes in the neuronal firing rate, as observed in our TIMM50 KD neurons, might occur in TIMM50 patients, and could lead to the epileptic phenotype seen in these patients. However, more studies are needed to verify this hypothesis.

In summary, our results suggest that even low levels of TIMM50 and TIM23^CORE^ components suffice in maintaining the majority of mitochondrial matrix and inner membrane proteome. Nevertheless, reduction in TIMM50 levels leads to a decrease of many OXPHOS and MRP complex subunits, which indicates that normal TIMM50 levels might be mainly essential for maintaining the steady state levels and assembly of intricate complex proteins. The consequently reduced cellular ATP levels and the detected mitochondrial abnormalities in neurons provide a plausible link between the TIMM50 mutation and the observed developmental defects in the patients. Moreover, the increased electrical activity resulting from decreased steady state levels of KCNA2 and KCNJ10 potassium channels plausibly link the TIMM50 mutation to the epileptic phenotype of patients, thus, providing a new direction for therapeutic efforts.

## Materials and methods

### Generation of primary human fibroblasts

Four mm punch biopsy samples were obtained from two TIMM50 patients (Patient 1 (P1) and Patient 2 (P2) carrying the mutation c.446C>T; p.Thr149Met and a normal family member that served as a healthy control (HC). Primary fibroblast cells were generated using standard procedures (50). In brief, each biopsy sample was cut into 12-15 pieces and 2-3 pieces were placed in six-well plate wells containing complete Dulbecco’s modified Eagle’s medium (DMEM; DMEM, 20% fetal bovine serum (FBS), 1% sodium pyruvate, 1% penicillin-streptomycin) and previously coated with 0.1% gelatin. The fibroblasts were grown for 2-3 weeks and passaged into 10 cm plates. Cells obtained from the first three passages were frozen and stored for further use. Following the generation of the primary fibroblast cells, genomic DNA purification and sequencing were performed to verify the presence of the mutation. Sequencing primers are found in S1 Table.

### Generation of TIMM50 knockdown (KD) mice primary cortical neuronal cultures

Mouse primary cortical neurons were harvested from P_0_/P_1_ pups and cultured using a previously described procedure (51). Plates were pre-coated with Matrigel (Corning, 354234) (diluted 1:1000 in Hank’s balanced salt solution (Satorius, 02-018-1A) with 10 mM HEPES, pH 7.4 (Fisher bioreagents, BP310-500)). For TIMM50 knockdown, three targeting shRNA sequences (Sh1, Sh2 and Sh3) and a scrambled (Scr) control sequence were designed. All shRNA sequences were cloned into the third-generation lentiviral vector pLL3.7 for expression under control of the U6 promotor. The same plasmid also encoded EGFP under control of the hSyn promotor, which allowed us to visualize and differentiate neuronal cells from other cells in the culture. To produce lentiviral particles, HEK293T/17 (ATCC, CRL-11268) cells were co-transfected with each of the designed shRNA vectors, together with the lentiviral helper constructs pMDLg-pRRE, pRSV-REV and CMV-VSVG, via calcium phosphate transfection. To generate TIMM50 KD neurons, the neuronal cultures were transduced on 4 days *in vitro* (DIV) with the generated lentiviruses and grown until 18 DIV. Immunoblotting with anti-TIMM50 antibodies were used to assess KD efficiency. shRNA oligo sequences and pLL3.7 sequencing primers are found in S1 Table.

### Immunoblotting

Fibroblasts were grown to ∼90% confluency on 10 cm cell culture plates, harvested using trypsin, washed twice with PBS, and then lysed using 50 μl of solubilization buffer (50 mM Na-HEPES, pH 7.4, 150 mM NaCl, 1.5 mM MgCl_2_, 10% glycerol, 1% Triton X-100, 1 mM EDTA, 1 mM EGTA supplemented with 400 μM of PMSF and 1:1000 dilution of protease inhibitor cocktail (GenDEPOT, P3200-020). Neurons (∼1×10^6^ cells) were cultured in the wells of a six-well plate, transduced on 4 DIV, grown until 18 DIV, and lysed by adding 50 μl of solubilization buffer to each plate well, following by scraping with a cell scraper. The protein concentration of both lyzed cultures was measured using Bradford reagent (BioRad, 500-0006) and appropriate protein amounts (20-100 μg) were loaded onto homemade polyacrylamide gels (12/14/16%) and separated using sodium dodecyl sulfate-polyacrylamide gel electrophoresis. The separated proteins were then transferred to a PVDF membrane (Millipore, IPVH00010) and immunodetection was carried out using antibodies against the target proteins. The list of antibodies used can be found in S2 Table. For fibroblasts, actin or GAPDH served as a loading control. For neurons, tubulin was used as loading control. ImageJ was used for densitometry analysis of protein expression levels. At least three biological repeats were performed for each immunoblotted protein (supplemental material S1 and S2 Raw images).

### Quantitative protein assessment and analysis

Label-free quantitative mass spectrometry was performed based on a published procedure (52). Spectra were searched against the Uniprot/Swiss-Prot mouse database (17,041 target sequences) for neuronal cells or the Uniprot/Swiss-Prot human database (20,379 target sequences) for fibroblast cells using the Andromeda search engine integrated into MaxQuant. Methionine oxidation (+15.9949 Da), asparagine and glutamine deamidation (+0.9840 Da), and protein N-terminal acetylation (+42.0106 Da) were variable modifications (up to 5 allowed per peptide), while cysteine was assigned a fixed carbamidomethyl modification (+57.0215 Da). Trypsin-cleaved peptides with up to two missed cleavages were considered in the database search. A precursor mass tolerance of ±20 ppm was applied prior to mass accuracy calibration and ±4.5 ppm after internal MaxQuant calibration. Other search settings included a maximum peptide mass of 6,000 Da, a minimum peptide length of 6 residues, 0.05 Da tolerance for Orbitrap and 0.6 Da tolerance for ion trap MS/MS scans. The false discovery rates for peptide spectral matches, proteins, and site decoy fractions were all set to 1 percent. Quantification settings were as follows: Re-quantification in a second peak finding attempt after protein identification; matched MS1 peaks between runs; a 0.7 min retention time match window after an alignment function was found with a 20-minute RT search space. Protein quantitation was performed using summed peptide intensities provided by MaxQuant. The quantitation method only considered razor plus unique peptides for protein level quantitation.

Data were obtained from three biological repeats, each involving three technical repeats (i.e., nine samples in total) for every cell type (neurons: Sh2 and Scr control; fibroblasts: P1, P2 and HC). Statistical analysis was performed using Perseus software. Protein levels were considered to be increased or decreased in TIMM50-deficient cells if they were significantly different (p-value < 0.05) and had a fold-change of at least 1.414, relative to what was measured in control cells. Mitochondrial protein classification was performed manually by comparing the obtained data with the MitoCarta3.0 human and mouse databases (53). Go term analysis was performed using the database for annotation, visualization and integrated discovery (DAVID) (54).

### Seahorse XF mito-stress and glycolysis stress tests

Fibroblasts were plated in Seahorse XF 96-well plates (Agilent, 103775-100) at 20-30% confluency, with experiments being carried out at ∼90% confluency. For neurons, ∼1×10^5^ cells were plated in each well of Seahorse XF 96-well plates, transduced with the Scr control or Sh2 constructs on 4 DIV and the experiment was carried out on 18 DIV. One-two hours prior to the experiment, the fibroblasts or neuronal cultures were washed and the medium was replaced with Seahorse XF DMEM, pH 7.4 (Agilent, 103575-100).

For the mito-stress test, the medium was supplemented with 1 mM sodium pyruvate (Sigma, S8636-100ML), 10 mM glucose (Merck, 1.08337.1000) and 2 mM glutamine (Biological Industries, 03-020-1B). The plates were loaded with oligomycin (Sigma-Aldrich, O4876), carbonyl cyanide-p-trifluoromethoxyphenylhydrazone (FCCP, Sigma-Aldrich, C2920), and rotenone (Sigma-Aldrich, R8875) together with antimycin A (Sigma-Aldrich, A8674), at final well concentrations of 1, 2, 0.5 and 0.5 μM, respectively. For the glycolysis stress test, the medium was only supplemented with glutamine. The plates were loaded with glucose, oligomycin and 2-deoxy-D-glucose (2-DG, Sigma-Aldrich, D8375-1G)) at final well concentrations of 10 mM, 1 μM and 50 mM, respectively. Plates were then loaded into the Seahorse XFe96 Extracellular Flux Analyzer and the experiments were carried using the manufacturer’s protocol. To normalize the oxygen consumption rate (OCR) or extracellular acidification rate (ECAR) values, fibroblasts were dyed with SynaptoGreen (Biotium, 70022) immediately at the end of each experiment and fluorescence levels were measured using a plate reader (BioTek Synergy HTX). In the case of neurons, the cells were dyed with DRAQ5 (BioLegend, 424101) immediately at the end of each experiment and imaged on an Incucyte SX5 live cell imaging and analysis system. Dyed nuclei were counted using the ImageJ particle analysis function. Three to six biological repeats, each involving three to six technical repeats, were performed for each experimental and control group.

### Mitochondrial DNA content

Total DNA was isolated form near-confluent fibroblasts or 18 DIV transduced neuronal cultures using a GeneElute Mammalian Genomic DNA Miniprep kit (Sigma, G1N350-1KT). Quantitative real-time PCR (StepOnePlus Real-Time PCR System) was then performed on each sample in the presence of SYBR green (PCR Biosystems, PB20.16-05). Expression levels were determined using the comparative cycle threshold (2^−ΔΔCt^) method, with the hypoxanthine guanine phosphoribosyl transferase (HPRT)-encoding gene serving as housekeeping gene. Primer sequences used are listed in S1 Table.

### Mitochondrial trafficking

Neuronal cultures were plated in a similar manner as described for membrane potential measurements. Neuronal cultures (4 DIV) were co-transfected with the TIMM50 KD or control plasmids, as well as a mito-dsRed-expressing plasmid (55), using 0.6 µg of each plasmid and 0.6 µl of Lipofectamine 2000 (Invitrogen, 11668-027). A transfection rate of 2-5% was seen the next day. This sparse transfection allowed for visualization of single neurons and their mitochondria, given the dramatically reduced background that allowed for tracking of mitochondrial movement in individual neurites. Following transfection, 10 DIV cultures were live-imaged with an iMIC inverted microscope equipped with a Polychrome V system (TILL photonics) and an ANDOR iXon DU 888D EMCCD camera (Andor, Belfast, Northern Ireland). Cells were imaged using a 60x oil immersion objective (Olympus), under temperature and CO_2_ control. Fields of view containing neurites stretching for at least 50 μm and not more than 100 μm from the cell body were imaged for 5 minutes, with a 3 second interval between each image. Individual neurites in each image set were selected using the segmented line tool in ImageJ. The images were then calibrated in time and space and turned into kymographs using the KymoToolBox ImageJ plugin (56). Each individual mitochondrion present in the neurite was then manually tracked on the kymograph using the segmented line tool to extract various parameters. Mitochondria were defined as static if they moved at a speed lower than 0.02 μm/sec.

### Determination of cellular ATP levels

Neuronal cells were plated into a 96-well cell culture plate at a density of 5×10^4^ cells per well. Cells were transduced with the Sh2 or Scr control constructs on 4 DIV and cellular ATP measurements were performed on 18 DIV using a Luminescent ATP Detection Assay Kit (Abcam, ab113849). To block luminescence signal contamination from adjacent wells, the lysis step of the assay was performed in the clear cell culture plate and the lysates were transferred to a white, flat bottom 96-well plate. Luminescence signals were read using the GloMax Navigator System.

### Intrinsic neuronal excitability and spontaneous activity

Neurons were plated at a density of 1.5×10^5^ cells/well of a 12-well plate and transduced on 4 DIV, with experiments being carried out on 16-20 DIV. Conventional whole cell recordings were performed with borosilicate thin wall glass capillaries (World Precision Instruments, TW150-3) with an input resistance of 4-5 MΩ. Series resistance ranged from 8-25 MΩ. An EPC-9 patch clamp amplifier was used in conjunction with PatchMaster software (HEKA Electronik, Lambrecht, Germany). The external solution consisted of 140 mM NaCl, 3 mM KCl, 2 mM CaCl_2_, 1 mM MgCl_2_, 10 mM HEPES, supplemented with 2 mg/ml glucose, pH 7.4, osmolarity adjusted to 305 mOsm. The internal solution consisted of 110 mM K gluconate, 10 mM KCl, 2 mM MgCl_2_, 10 mM HEPES, 10 mM glucose, 10 mM Na creatine phosphate, and 10 mM EGTA, pH 7.4. Osmolarity adjusted to 285 mOsm.

To measure the rheobase and assess repetitive and maximal firing, long (250 ms) depolarization square current steps of varying intensity (from −100 to 600 pA, at 2 pA intervals) were applied. Signals were filtered at 2 kHz and sampled at 5 kHz. For α-dendrotoxin (α-DTX) measurements, long (250 ms) depolarizing square current steps of varying intensity (From 0-500 pA, at 100 pA intervals) were applied to the cells, before and after application of an external solution containing 100 nM α-DTX (Alomone Labs, D-350). Clear traces in the range of 100-300 pA were chosen in the before-α-DTX measurements and compared to the respective traces in the after-α-DTX measurements. To measure spontaneous activity, an external solution containing 1 μM tetrodotoxin (TTX; Alomone Labs, T550_1mg) was perfused onto the cells, followed by a high current injection to assure that no action potentials were being generated. The mode was then switched to voltage clamp, the holding voltage was adjusted to −60 mV and mEPSCs were recorded for 2 minutes (at repeating 10 second intervals). The data were analyzed with Igor Pro software (Wavemetrics, Lake Oswego, OR). The TaroTools procedure set (for Igor Pro) was used to analyze spontaneous activity measurements.

## Supporting information

Supplemental raw images 1

Supplemental raw images 2

## Acknowledgements

The authors thank Professor Bernard Attali, Dr. Celeste Weiss Katz, Dr. Natalia Borovok and Dr. Amit Kessel for their valuable input and guidance, and to the undergrad project students Afek Moravia and Avia Tamir for their technical support. This work was supported by the Emory University Emory Integrated Proteomics Core Facility (RRID:SCR_023530). A.A. is incumbent of The Louise and Nahum Barag Chair in Molecular Genetics of Cancer. U.A. is the incumbent of the Michael Gluck Chair in Neuropharmacology and A.L.S Research.

## Author contributions

**Eyal Paz**: Writing-original draft; data curation; formal analysis; investigation. **Sahil Jain:** Writing-original draft; data curation; formal analysis; investigation. **Irit Gottfried:** resources; writing-review and editing; validation; **Orna Staretz-Chacham**: Resources. **Muhammad Mahajnah**: Resources. **Pritha Bagchi:** Investigation; writing-review and editing. **Nicholas T. Seyfried:** Supervision; methodology. **Uri Ashery:** Conceptualization; supervision; funding acquisition; validation; methodology; writing-review and editing. **Abdussalam Azem:** Conceptualization; supervision; funding acquisition; validation; methodology; writing-review and editing.

## Supporting information

**Fig S1. PCA analysis for fibroblasts proteomics replicates.**

PCA analysis showing that the proteomics P1 and P2 replicates are consistently different than the HC replicates.

**Fig S2. Immunoblot confirmation of mass spectrometry-based proteomics findings.**

(A) Verification of the measured increase in the levels of ALDH2 and GRSF1 in fibroblasts. Three biological repeats were performed for each antibody. Full results and original blots are found in supplemental material S1 Raw images. (B) Verification of the measured decrease in the levels of the KCNA2 potassium channel in neuronal cultures. Three biological repeats were performed. Full results and original blots are found in supplemental material S2 Raw images.

**Fig S3. Go term analysis of all significantly decreased fibroblast proteins and OXA1 immunoblot.**

(A) Go term analysis confirms the enrichment of inner mitochondrial membrane proteins, OXPHOS related processes and mitochondrial ribosome subunits. (B) Healthy control- and patients-derived primary fibroblasts were lysed and analyzed by immunoblot with OXA1 antibody. Actin was used as loading control. (C) Band density analysis shows no significant change in the levels of OXA1. Band signals were normalized to the loading control and compared to the level measured in the healthy control sample, taken as 100%. Data are shown as means ± SEM, n = 3 biological repeats. Full results and original blots are found in supplemental material S1 Raw images.

**Fig S4. Spontaneous excitatory activity recordings and the effect of α-DTX on TIMM50 KD neuronal cells.**

(A) Representative traces of mEPSCs measurements taken in the presence of 1 μM TTX. (B) No change in the average mEPSC amplitude, frequency and area under the peak was observed in TIMM50 KD neuronal cells compared to Scr control-transduced cells. Data are shown as means ± SEM, n = 23 / 20 cells (for Scr control-/Sh2-transduced cells, respectively) from three biological repeats, Mann-Whitney test. (C) Histogram of the relative frequency distribution of the mEPSC amplitude measurements. (D) Cumulative distribution function of the mEPSC amplitude measurements. (E) Measurements of the number of action potentials fired before and after treatment with 100 nM α-DTX for Scr control- and Sh2-transduced neuronal cells. (F) A higher difference in the number of action potentials fired after and before α-DTX application was observed in Scr control-transduced cells compared to TIMM50 KD neuronal cells. n = 21 / 20 cells (for Scr control-/Sh2-transduced cells, respectively) from three biological repeats, ***p-value < 0.001, unpaired Student’s t-test.

**S1 Table. Oligonucleotides used in this study.**

**S2 Table. Reagents and tools used in this study.**

**S1 Raw images. Original blots – Fibroblasts.**

**S2 Raw images. Original blots – Neurons.**

**S1 Dataset. HC vs P1 all proteins fold change.**

**S2 Dataset. HC vs P2 all proteins fold change.**

**S3 Dataset. Scr vs Sh2 all proteins fold change.**

**S1 Movie. Representative mitochondrial trafficking movies.** Scr control neurons and TIMM50 KD neurons co-transfected with the corresponding control/KD plasmid and mito-DsRed plamid. The cells were imaged for 5 minutes with a 3 second interval between each image. The videos displayed are sped up by ∼15x.

