## Supplementary figures and images for "Biochemical and neurophysiological effects of deficiency of the mitochondrial import protein TIMM50"

### Supplemental figure 1

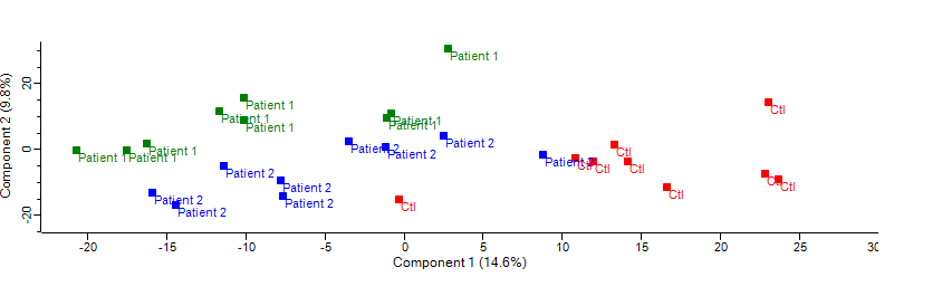

### Supplemental figure 2

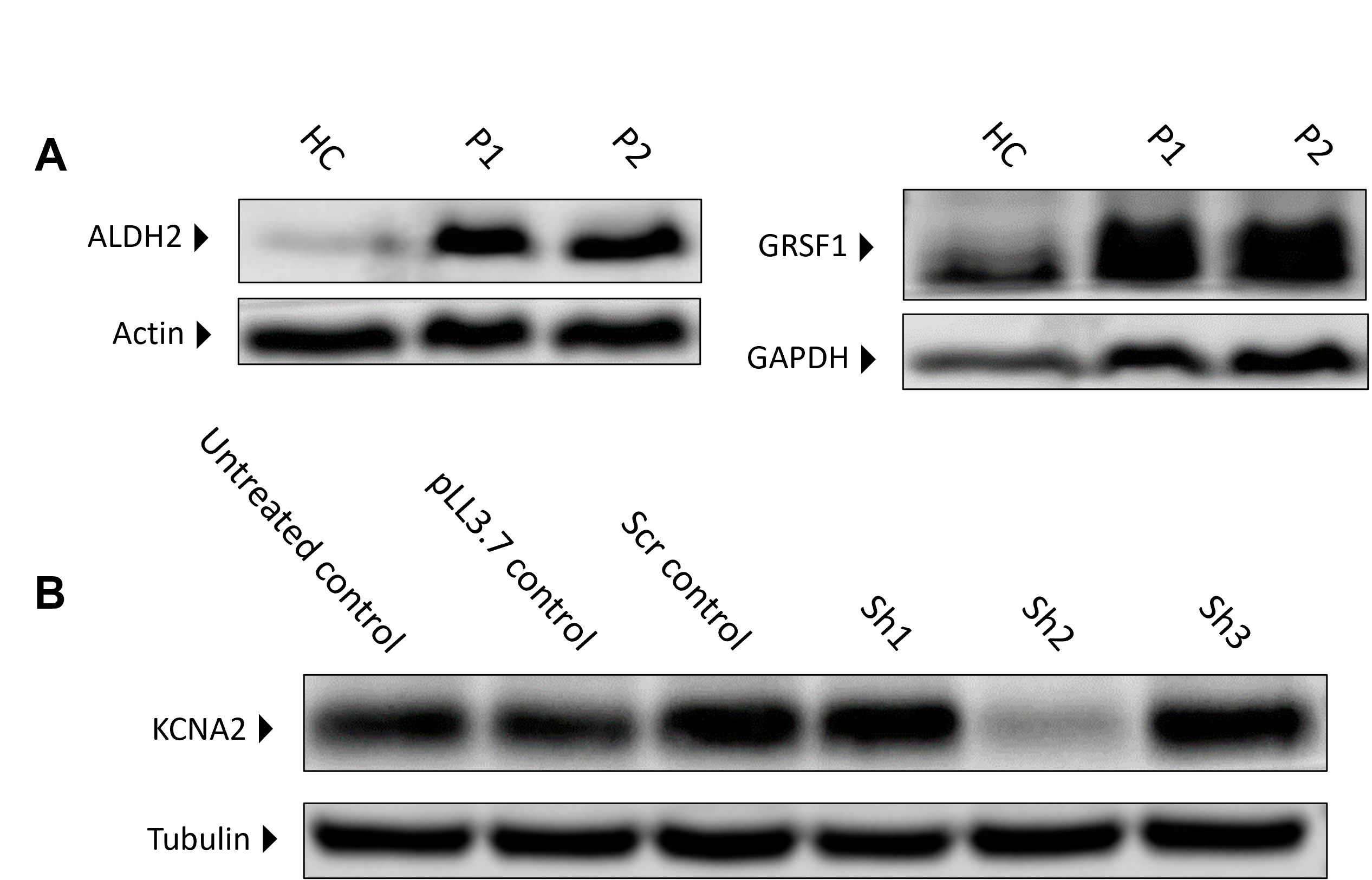

### Supplemental figure 3

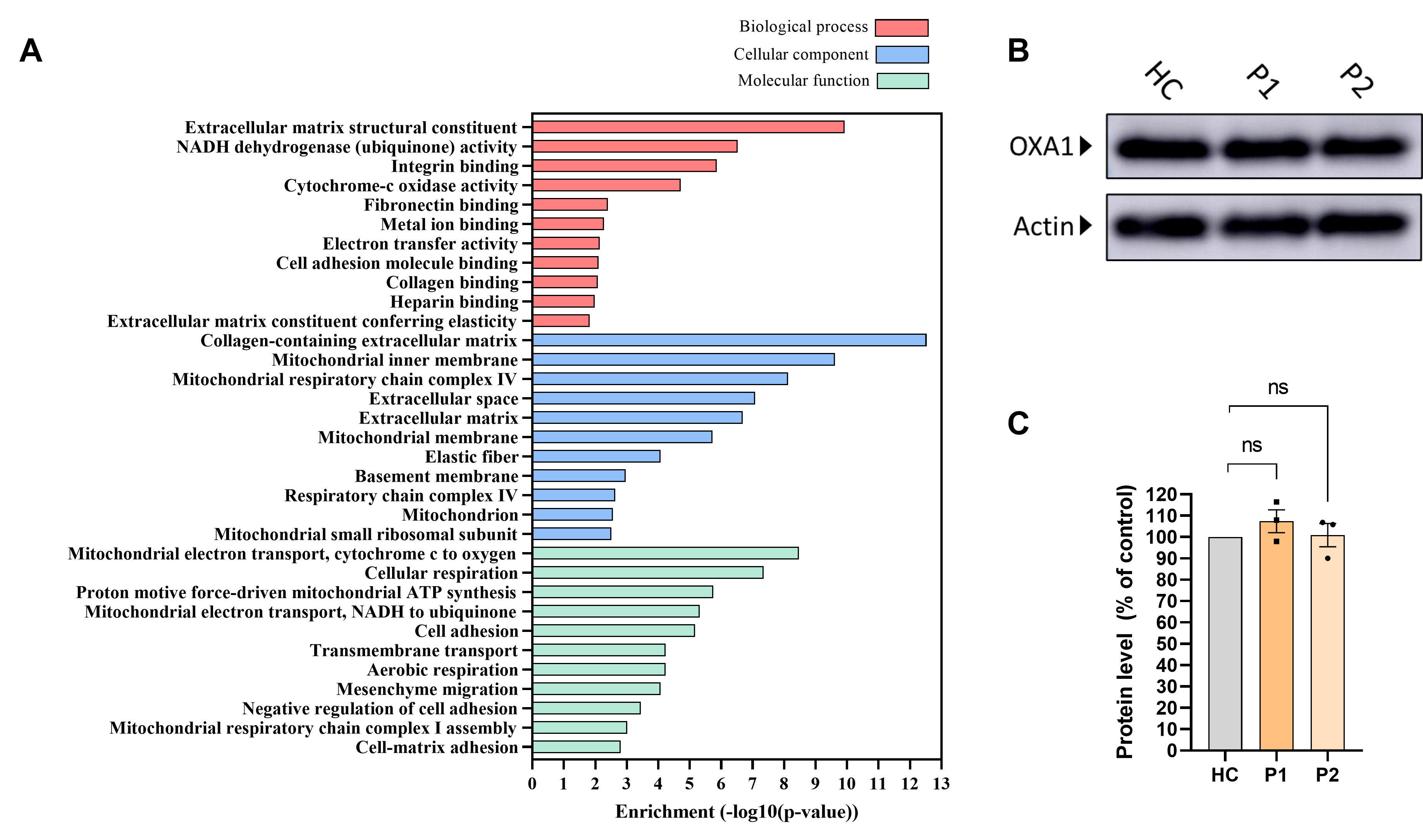

### Supplemental figure 4

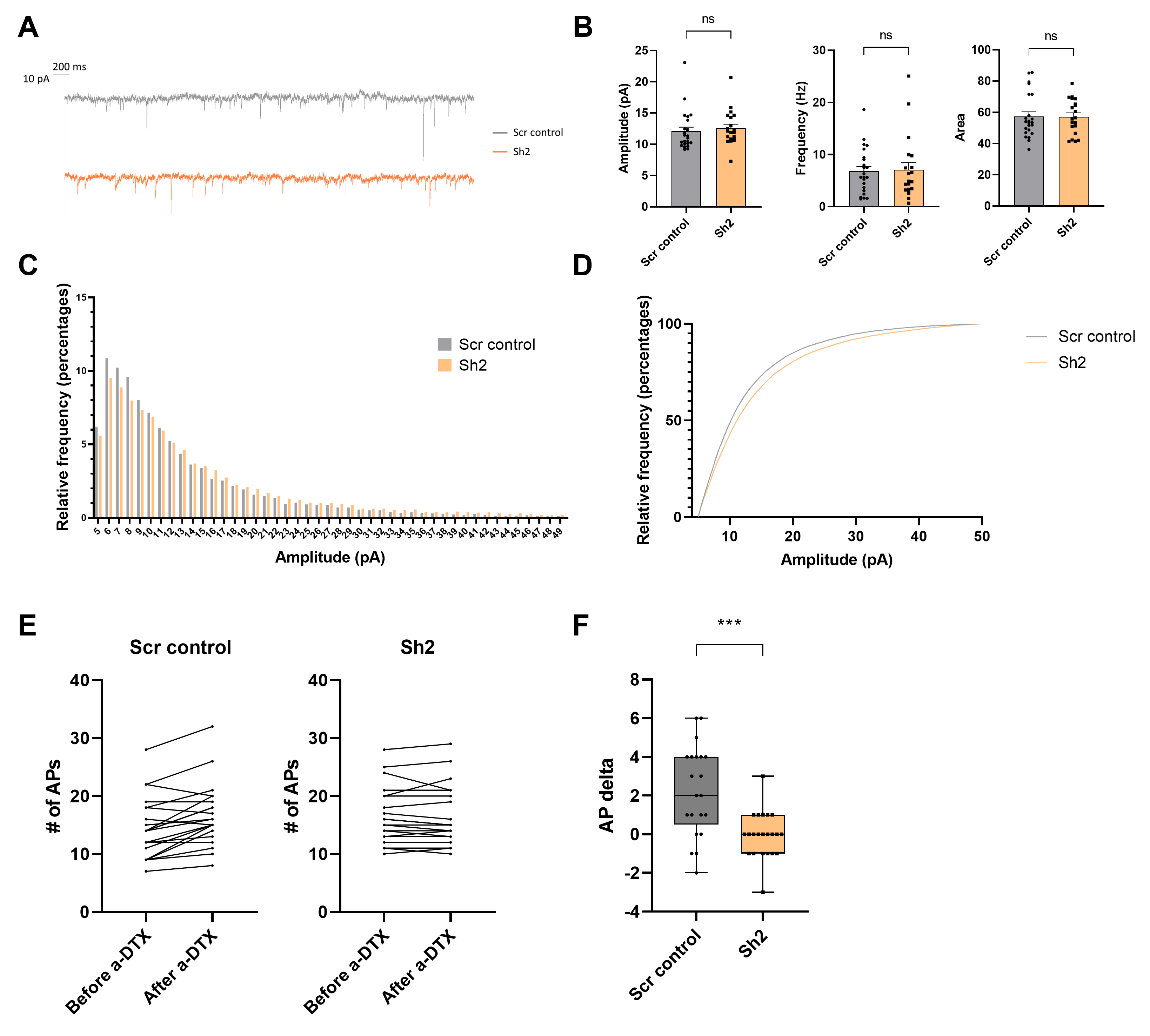

### Supplemental movie 1

#### Slide 1
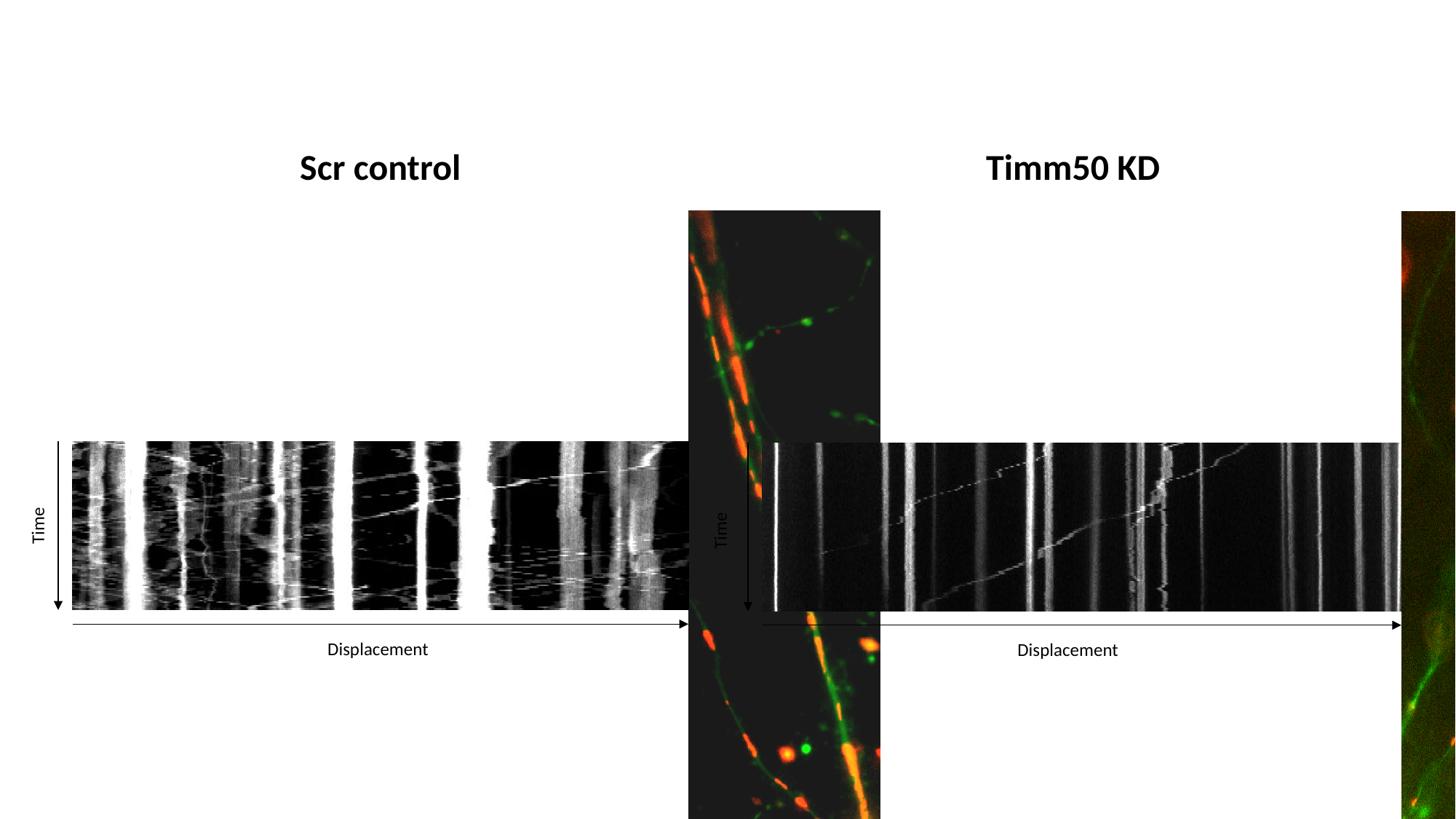

Scr control
Timm50 KD
Time
Time
Displacement
Displacement
