## Supplemental table 1 for "Biochemical and neurophysiological effects of deficiency of the mitochondrial import protein TIMM50"

| **Sequence** | **Application** |
| --- | --- |
| **Sequencing primers** | |
| 5'-GCATTCTGTTTCATACCTGG-3' | Forward primer for the human TIMM50 gene, to confirm the mutation presence. |
| 5'-GGCTGAAAGTCCACCTCATC-3' | Reverse primer for the human TIMM50 gene, to confirm the mutation presence. |
| 5'-GGAAACTCACCCTAACTGTAAAG-3' | Forward primer for pLL3.7 hSyn sequencing of shRNA segment. |
| 5'-GGTCCTAAAACCCACTTGCACTC-3' | Reverse primer for pLL3.7 hSyn sequencing of shRNA segment. |
| **qPCR primers** | |
| 5′-AATCTACCATCCTCCGTGAAACC-3′ | Forward primer for Dloop1 Gene |
| 5′-TCAGTTTAGCTACCCCCAAGTTTAA-3′ | Reverse primer for Dloop1 Gene |
| 5′-CTAGCTCATGTGTCAAGACCCTC-3′ | Forward primer for TERT Gene |
| 5′-GCCAGCACGTTTCTCTGTT-3′ | Reverse primer for TERT Gene |
| 5′-GCAGTACAGCCCCAAAATGG-3′ | Forward primer for HPRT Gene |
| 5′-GGTCCTTTTCACCAGCAAGCT-3′ | Reverse primer for HPRT Gene |
| **shRNA oligo sequences** | |
| 5′-AGCTGTCCACACTTTGCCG-3′ | Sh1 |
| 5′-AAGGACCCCGGTAAGCTCC-3′ | Sh2 |
| 5′-AAACCGGCGATCCAGAGCG-3′ | Sh3 |
| 5′-AAGGAAACCGTCGCGCCGA-3′ | Scr control |
