## Supplemental table 2 for "Biochemical and neurophysiological effects of deficiency of the mitochondrial import protein TIMM50"

| **Regent/Resource** | **Reference or Source** | **Identifier or Catalog Number** |
| --- | --- | --- |
| **Experimental models** |  |  |
| HEK293T/17 | ATCC | CRL-11268 |
| C57BL/6JRccHsd | Envigo | 1BL/6R18 |
| Patient 1 fibroblasts | Derived from a skin biopsy obtained by Dr. Muhammad Mahajnah and Dr. Orna Staretz-Chacham | N/A |
| Patient 2 fibroblasts | Derived from a skin biopsy obtained by Dr. Muhammad Mahajnah and Dr. Orna Staretz-Chacham | N/A |
| Healthy control fibroblasts | Derived from a skin biopsy obtained by Dr. Muhammad Mahajnah and Dr. Orna Staretz-Chacham | N/A |
| **Recombinant DNA** |  |  |
| pLL3.7-hSyn |  |  |
| pLL3.7-hSyn-Scr control | This study | N/A |
| pLL3.7-hSyn-Sh1 | This study | N/A |
| pLL3.7-hSyn-Sh2 | This study | N/A |
| pLL3.7-hSyn-Sh3 | This study | N/A |
| pMDLg-pRRE | Addgene | 12251 |
| pRSV-REV | Addgene | 12253 |
| pCMV-VSV-G | Addgene | 8454 |
| pDSRed-mito | Lindenboim et al (2013) (Cellular and Molecular Life Sciences) | Prof. Reuven Stein, TAU, Israel |
| **Antibodies** |  |  |
| Rabbit anti-TIMM50 (For fibroblasts) | Proteintech | 22229-1-AP |
| Rabbit anti-TIMM50 (For fibroblasts) | Abcam | ab204486 |
| Rabbit anti-TIMM50 (For neurons) | Abcam | ab109436 |
| Mouse anti-TIMM23 (For neurons) | Santa Cruz | sc-514463 |
| Rabbit anti-TIMM23 (For fibroblasts) | Abcam | ab230253 |
| Rabbit anti-TIMM17A | Abcam | ab192246 |
| Rabbit anti-TIMM17B | Abcam | ab234888 |
| Rabbit anti-TIMM21 | LSBio | LS-C803032 |
| Rabbit anti-TIMM44 (For fibroblasts) | Abcam | ab194829 |
| Rabbit anti-TIMM44 (For neurons) | Abcam | ab201453 |
| Rabbit anti-Pam16 | Abcam |  |
| Rabbit anti-TOMM40 (For fibroblasts) | Cell signaling | 55959 |
| Rabbit anti-TOMM40 (For neurons) | Abcam | ab272921 |
| Rabbit anti-TOMM20 | Cell signaling | 42406 |
| Rabbit anti-mtHsp60 | Prepared in house, raised against purified human mtHsp60 | ab228923 |
| Rabbit anti-Aconitase-2 | Abcam | ab228923 |
| Rabbit anti-ALDH2 | Abcam | ab227021 |
| Rabbit anti-GRSF1 | Abcam | ab205531-100 |
| Mouse anti-OXA1 | Abcam | ab88975 |
| Mouse anti-KCNA2 | Upstate Biotechnology | 05-408 |
| Mouse anti-α-Tubulin (Loading ctrl for neurons) | Sigma Aldrich | T5168 |
| Mouse anti-β-Actin (Loading ctrl for fibroblasts) | Sigma Aldrich | A1978 |
| Rabbit anti-GAPDH (Loading ctrl for fibroblasts) | Cell signaling | 5174 |
| Goat α Mouse | Jackson | 115-035-003 |
| Goat α Rabbit | Jackson | 111-035-144 |
| **Oligonucleotides and sequence-based reagents** |  |  |
| Sequencing primers | Hylabs | Dataset EV4 |
| qPCR primers | Hylabs | Dataset EV4 |
| shRNA oligo sequences | Hylabs | Dataset EV4 |
| **Chemicals, enzymes and other reagents** |  |  |
| DMEM | Satorius | 01-052-1A |
| Fetal bovine serum (FBS) | Gibco | 10270-106 |
| Sodium Pyruvate | Sigma | S8636-100ML |
| Penicillin-Streptomycin | Rhenium | 15140-122 |
| Neurobasal-A | Gibco | 10888-022 |
| B-27 | Gibco | 17504-044 |
| 5-Fluoro-2'-deoxyuridine | Sigma | F0503-100MG |
| GlutaMAX | Gibco | 35050-038 |
| Uridine | Sigma | U3750-25G |
| Papain | Sigma | P4762-100MG |
| DNase | Sigma | D5025-15KU |
| L-Cysteine | Sigma | C7352-25G |
| Hanks Balanced Salt Solution | Satorius | 02-018-1A |
| Matrigel | Corning | 354234 |
| HEPES | Fisher bioreagents | BP310-500 |
| NaCl | Emsure | 1.06404.1000 |
| MgCl2 | Acros organics | 447155000 |
| Glycerol | Bio-lab | 000712050100 |
| Triton X-100 | Sigma | T9284-1L |
| EDTA | J.T. Baker | 8993-01 |
| EGTA | Sigma | E0396-10G |
| Phenylmethylsulfonyl fluoride (PMSF) | Roche | 10 837 091 001 |
| protease inhibitor cocktail | GenDEPOT | P3200-020 |
| Bradford reagent | BIO-RAD | 500-0006 |
| PVDF membrane | Millipore | IPVH00010 |
| Seahorse XF 96 well plates | Agilent | 103775-100 |
| Seahorse XF DMEM Medium, pH=7.4 | Agilent | 103575-100 |
| Glucose | Merck | 1.08337.1000 |
| Glutamine | Biological Industries | 03-020-1B |
| Oligomycin | Sigma-Aldrich | O4876 |
| Carbonyl cyanide-p-trifluoromethoxyphenylhydrazone (FCCP) | Sigma-Aldrich | C2920 |
| Rotenone | Sigma-Aldrich | R8875 |
| Antimycin A | Sigma-Aldrich | A8674 |
| 2-Deoxy-D-glucose | Sigma-Aldrich | D8375-1G |
| SynaptoGreen | Biotium | 70022 |
| DRAQ5 | BioLegend | 424101 |
| ATP Detection Assay Kit | Abcam | ab113849 |
| GeneElute Mammalian Genomic DNA Miniprep kit | Sigma | G1N350-1KT |
| SYBR Green | PCRBIOSYSTEMS | PB20.16-05 |
| Lipofectamine 2000 | Invitrogen | 11668-027 |
| KCl | Emsure | 1.04936.1000 |
| CaCl2 | Acros organics | 207780010 |
| K Gluconate | Sigma | P1847-100G |
| Na Creatine-phosphate | Sigma | 27920-1G |
| Borosilicate thin wall glass capillaries | World Precision Instruments | TW150-3 |
| α-Dendrotoxin | Alomone Labs | D-350 |
| Tetrodotoxin (TTX) | Alomone Labs | T550_1mg |
| **Software** |  |  |
| Graphpad Prism 8.0.2 | https://www.graphpad.com/ |  |
| ImageJ | https://imagej.nih.gov/ij/index.html |  |
| Snapgene Viewer | https://www.snapgene.com/snapgene-viewer |  |
| Perseus | https://maxquant.net/perseus/ |  |
| PatchMaster software | https://www.heka.com/index.html |  |
| IGOR Pro v6.0.4.0 | https://www.wavemetrics.com/ |  |
| **Other** |  |  |
| Seahorse XFe96 Extracellular Flux Analyzer | Agilent |  |
| Synergy HTX | BioTek |  |
| Incucyte SX5 live cell imaging and analysis system | Creative Biolabs |  |
| GloMax Navigator System | Promega |  |
| StepOnePlus Real-Time PCR System | Thermo Fisher |  |
| Olympus IX81 microscope supplemented with Abberior STEDYCON device | Olympus, Abberior |  |
| iMIC inverted microscope equipped with Polychrome V system and an ANDOR iXon DU 888D EMCCD camera | TILL photonics, Andor |  |
| EPC-9 patch clamp amplifier | HEKA Electronik Gmbh |  |
