## Supplemental raw images 1 for "Biochemical and neurophysiological effects of deficiency of the mitochondrial import protein TIMM50"

### TIMM50

(The running order is H.C / P1 / P2)

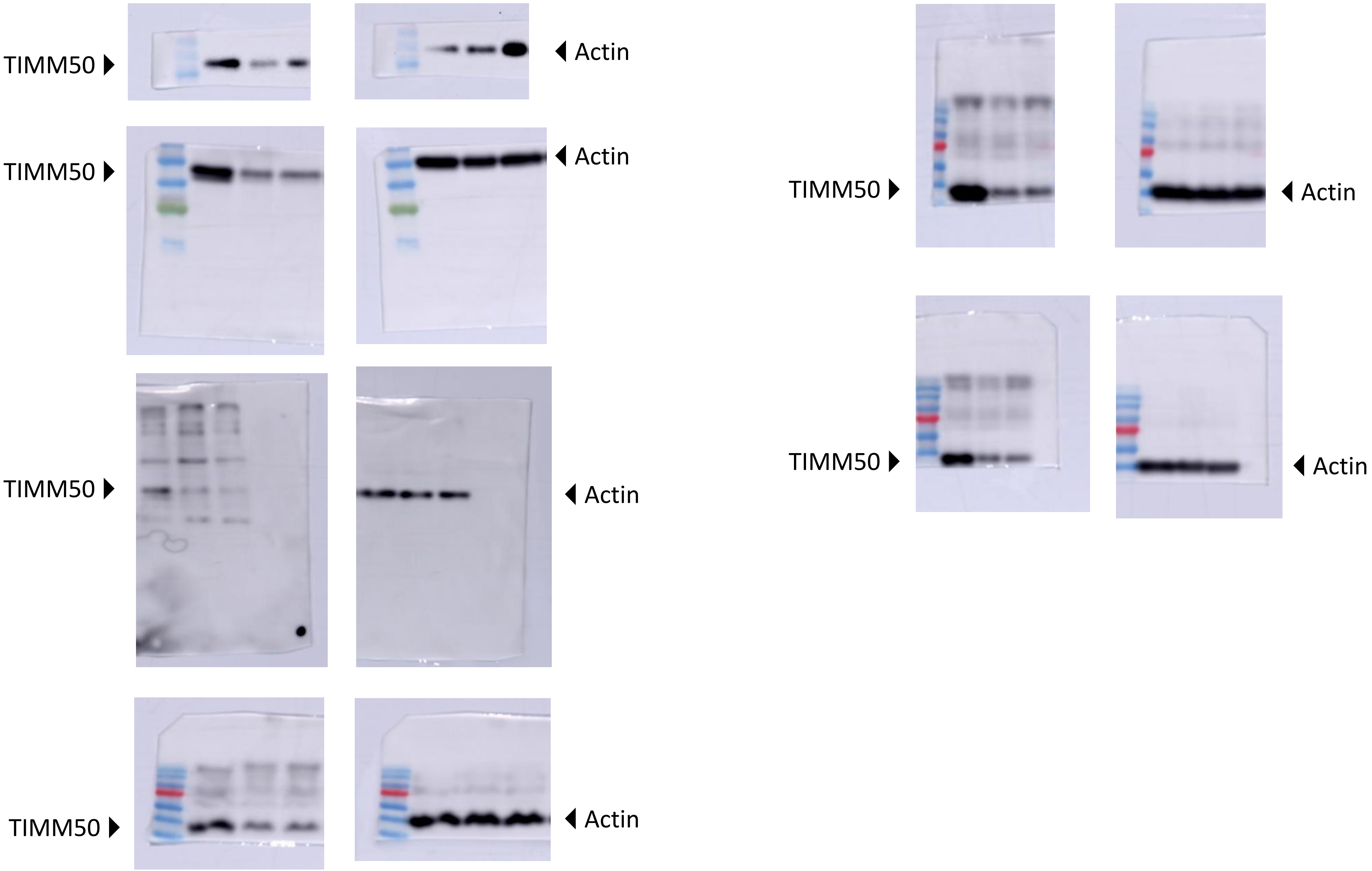

### TIMM23

(The running order is H.C / P1 / P2)

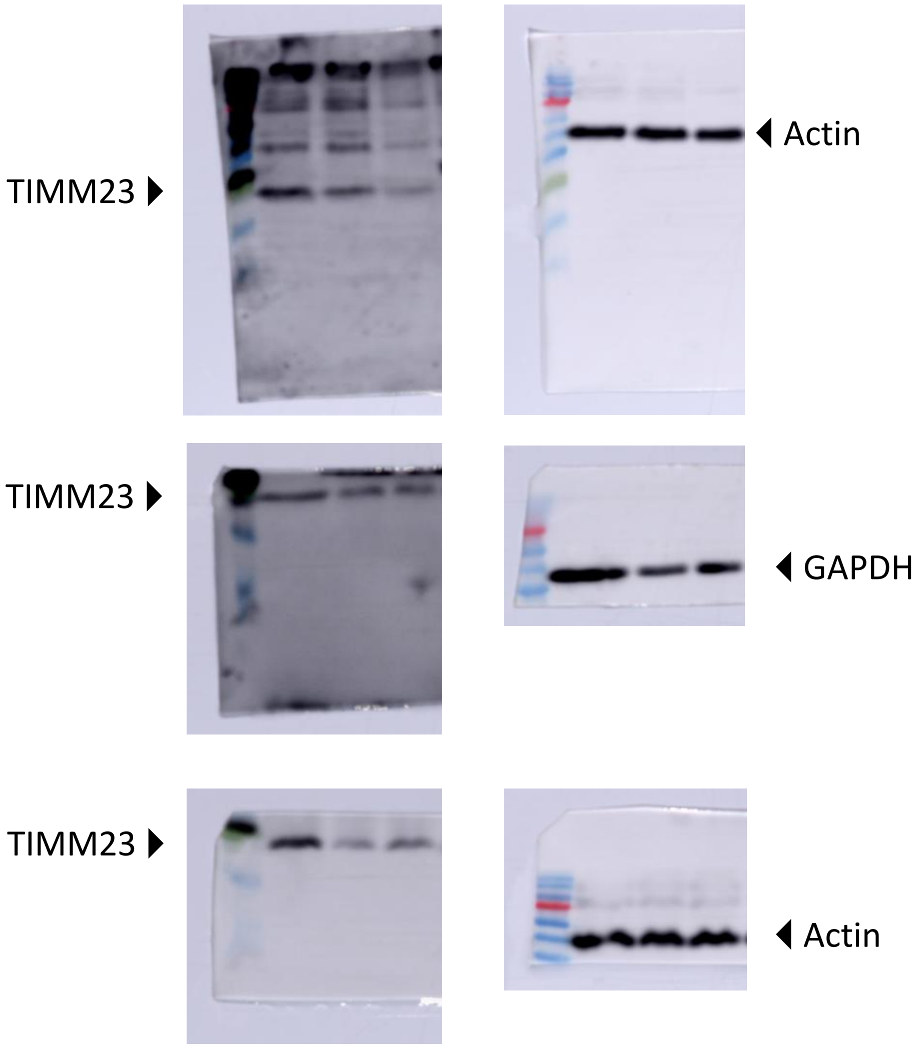

### TIMM17A

(The running order is H.C / P1 / P2)

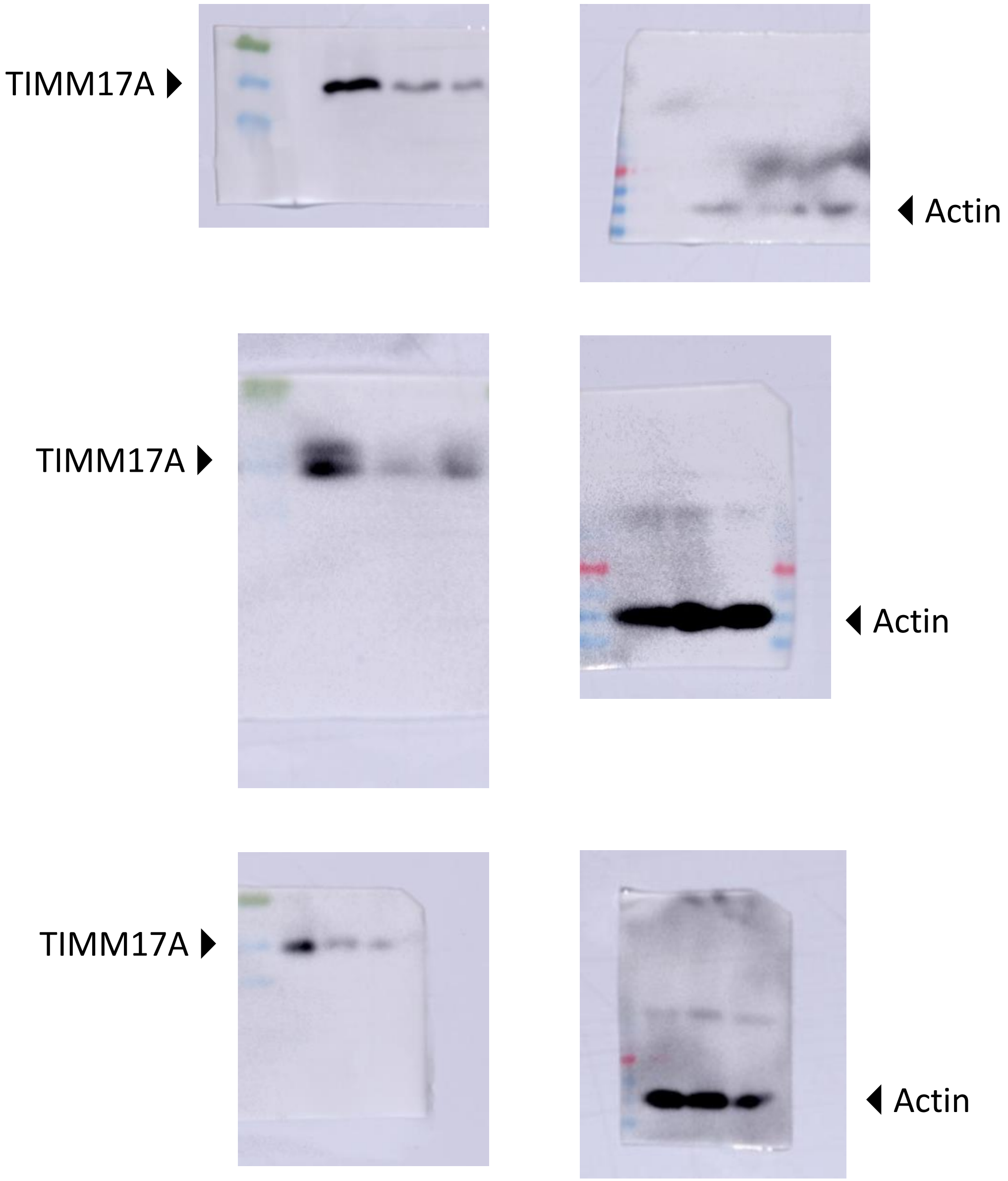

### TIMM17B

(The running order is H.C / P1 / P2)

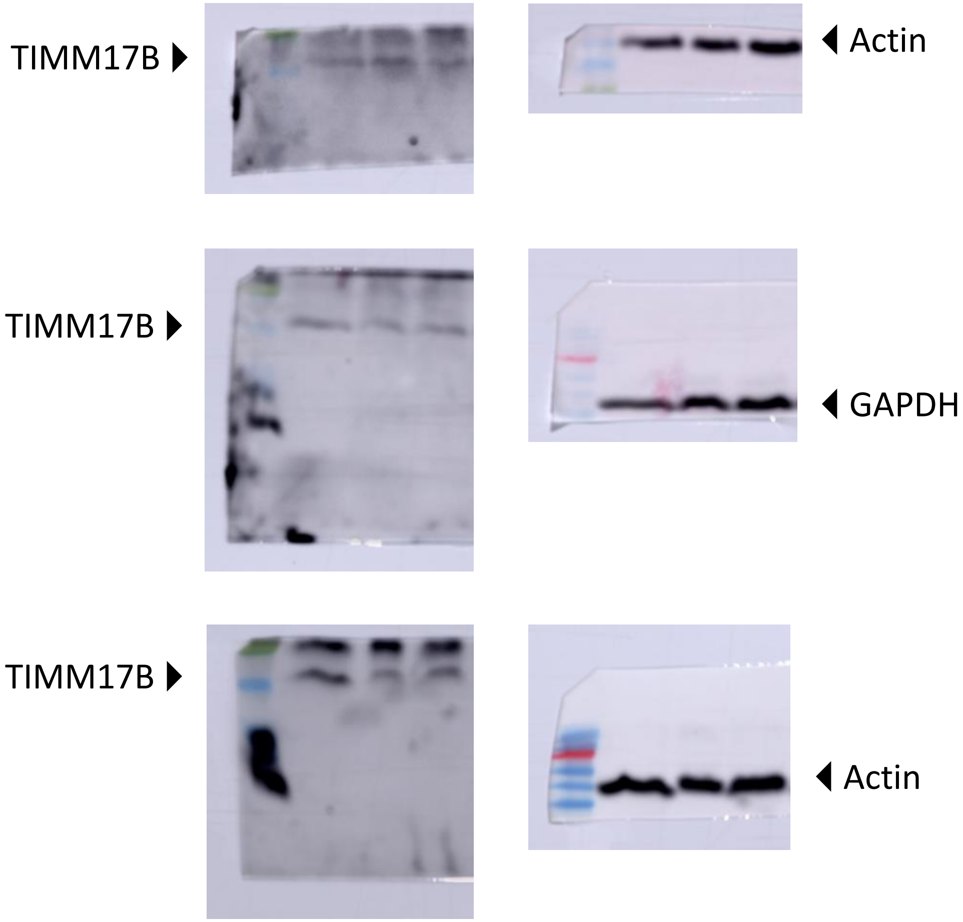

### TIMM21

(The running order is H.C / P1 / P2)

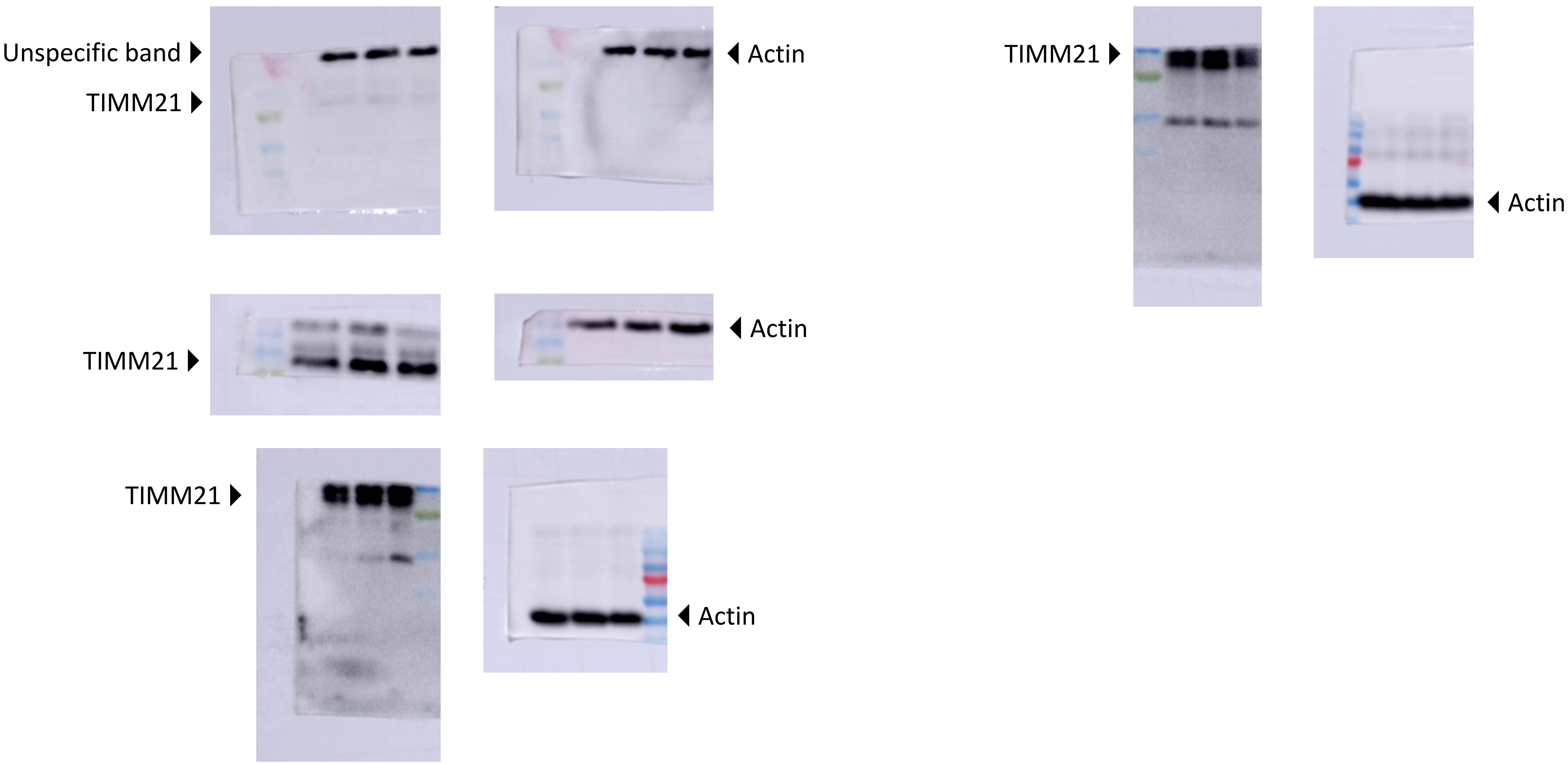

### TIMM44

(The running order is H.C / P1 / P2)

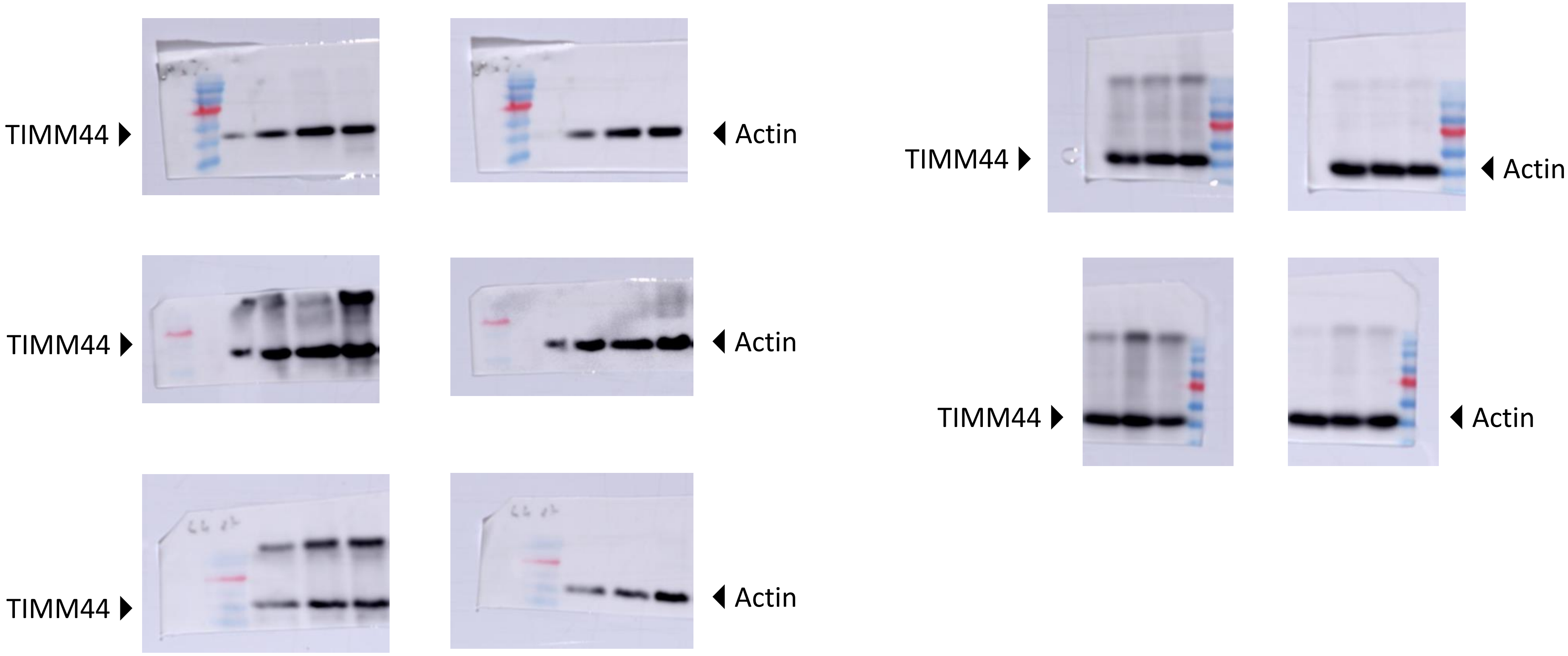

### Pam16

(The running order is H.C / P1 / P2)

Pam16 ▶
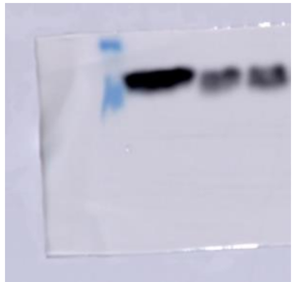
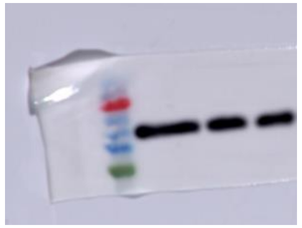
◀ Actin

Pam16 ▶
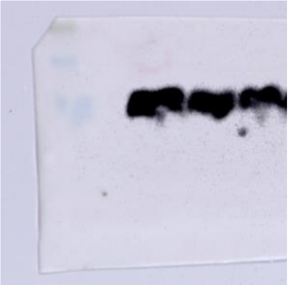
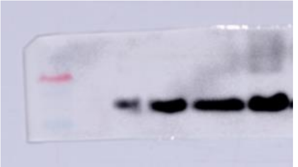
◀ Actin

Pam16 ▶
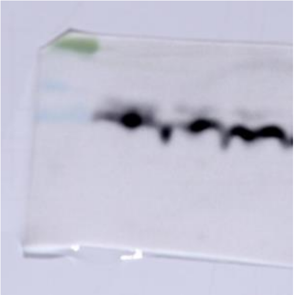
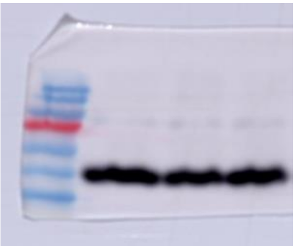
◀ Actin

### TOMM40

(The running order is H.C / P1 / P2)

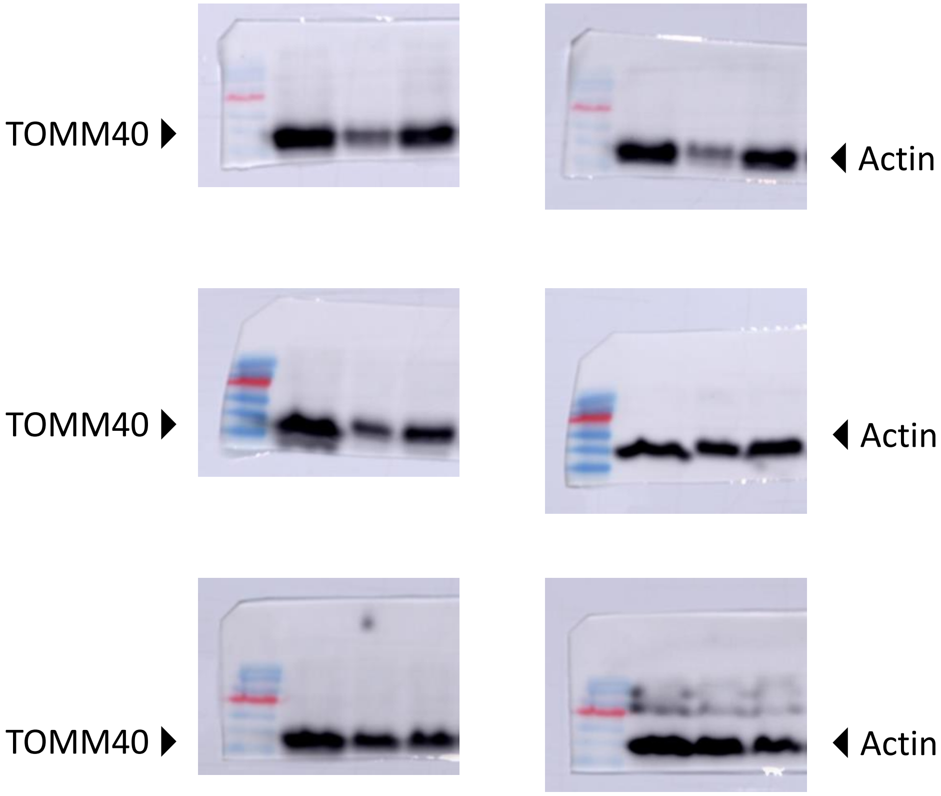

### TOMM20

(The running order is H.C / P1 / P2)

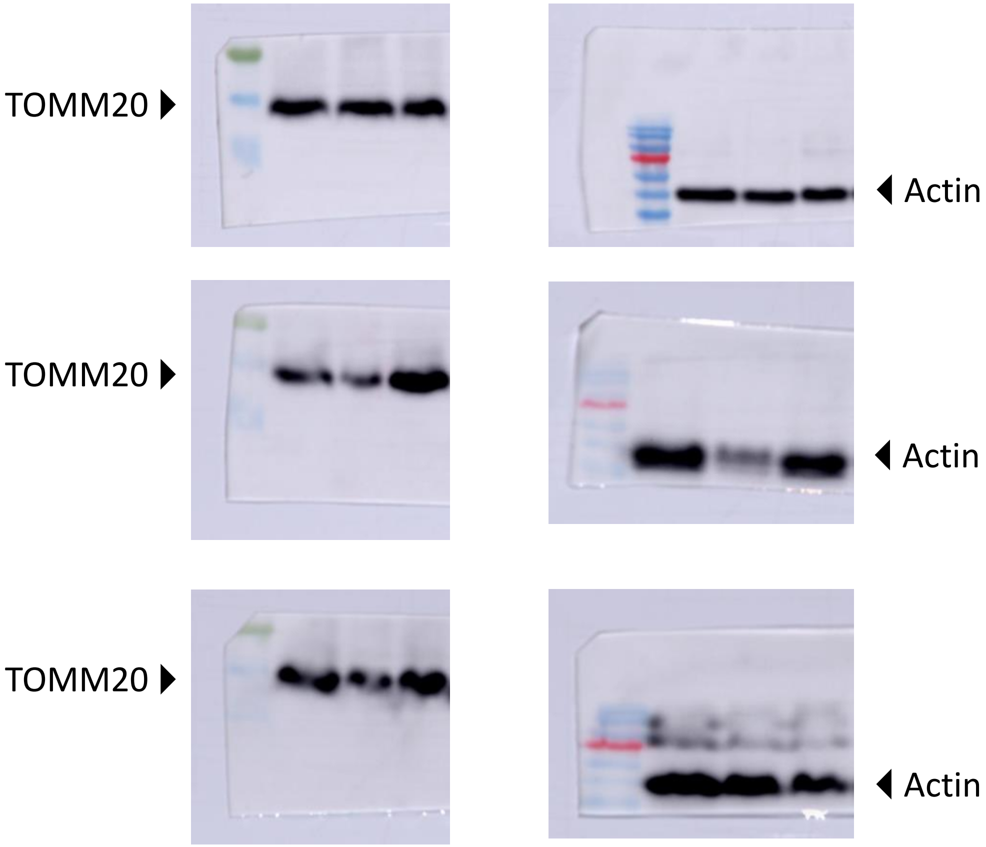

### mtHsp60

(The running order is H.C / P1 / P2)

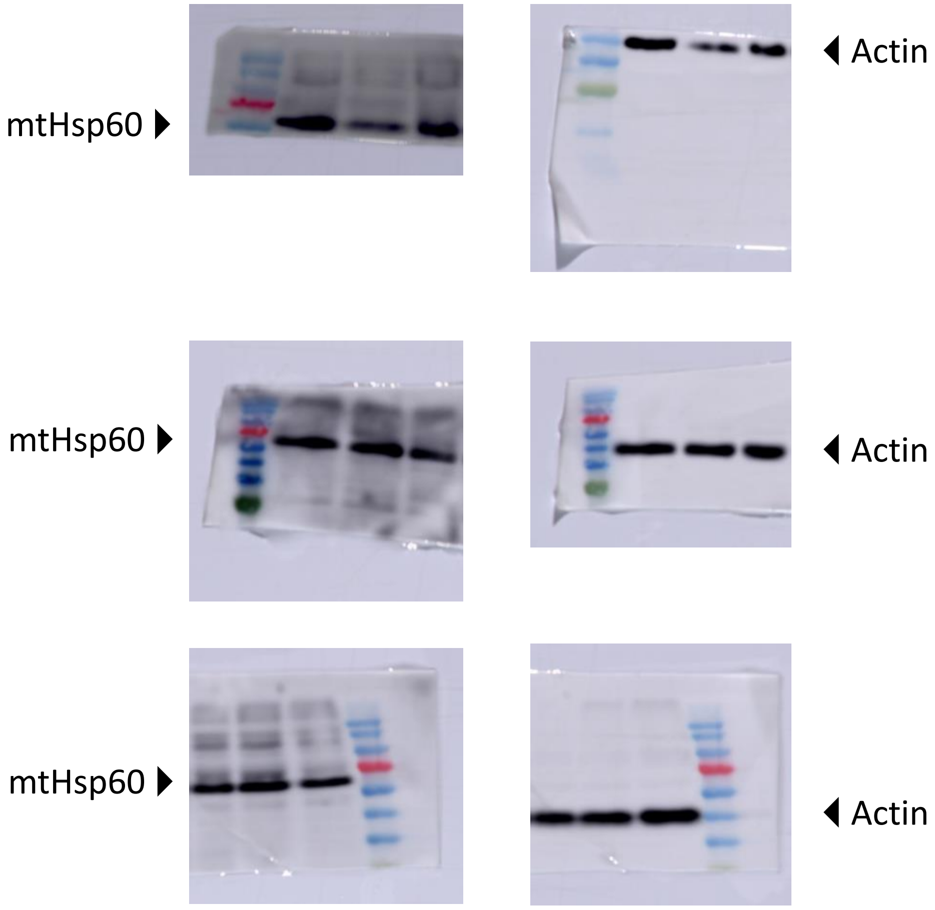

### Aconitase-2

(The running order is H.C / P1 / P2)

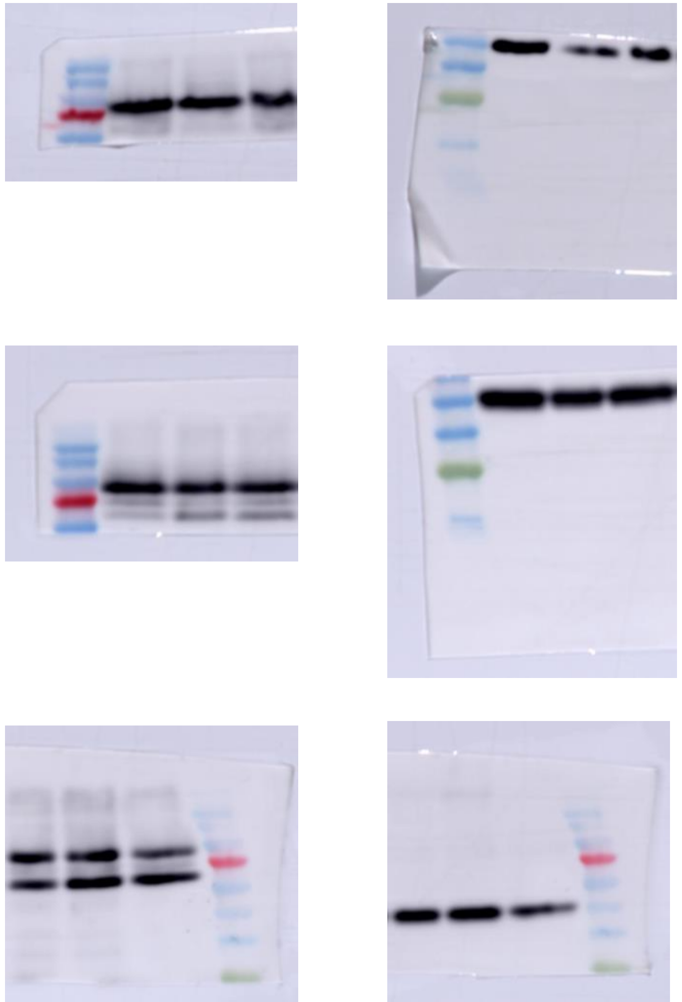

### ALDH2

(The running order is H.C / P1 / P2)

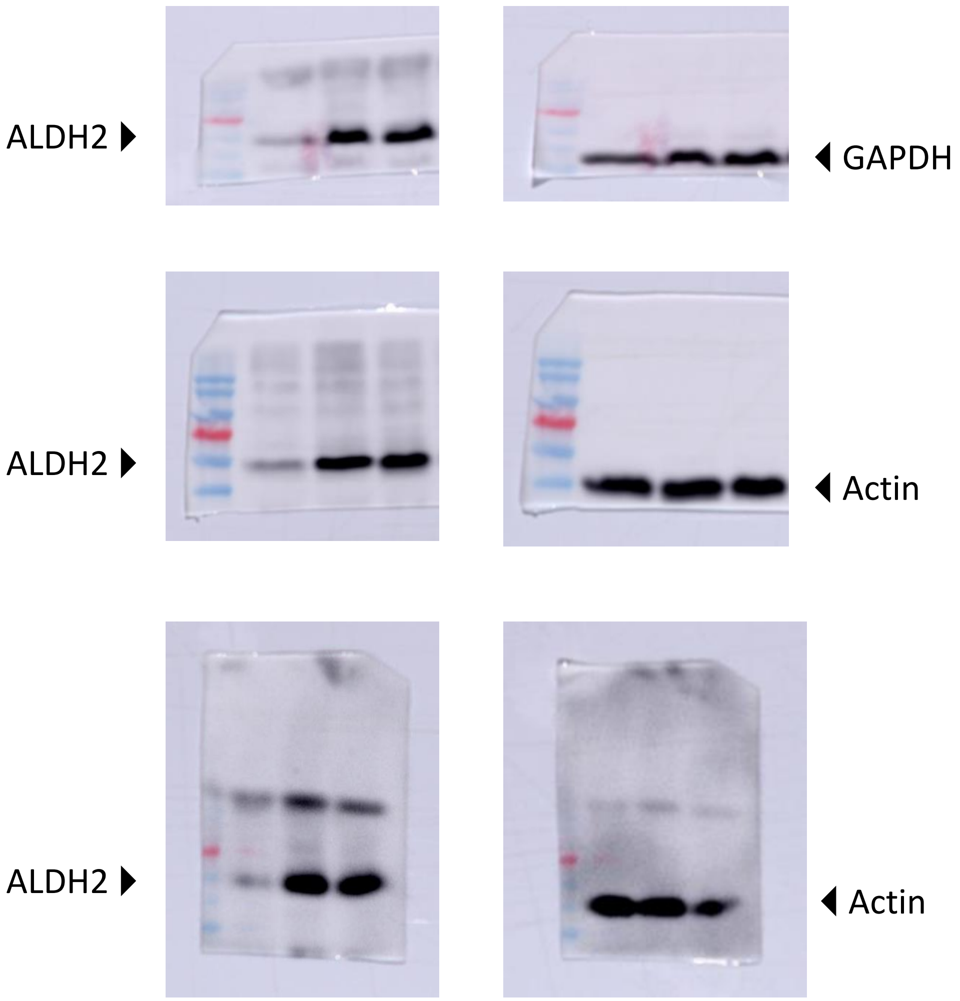

### GRSF1

(The running order is H.C / P1 / P2)

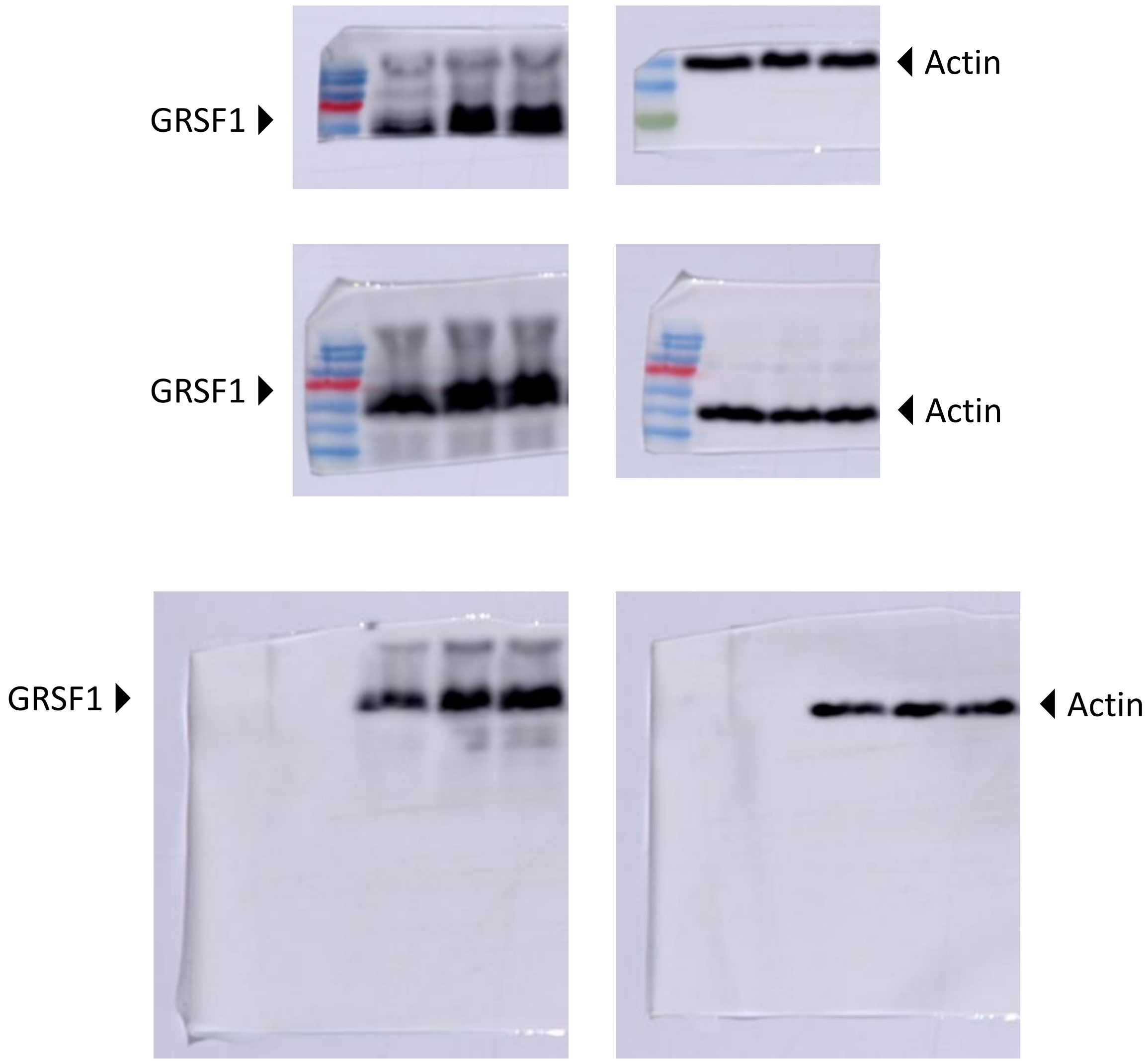

### OXA1

(The running order is H.C / P1 / P2)

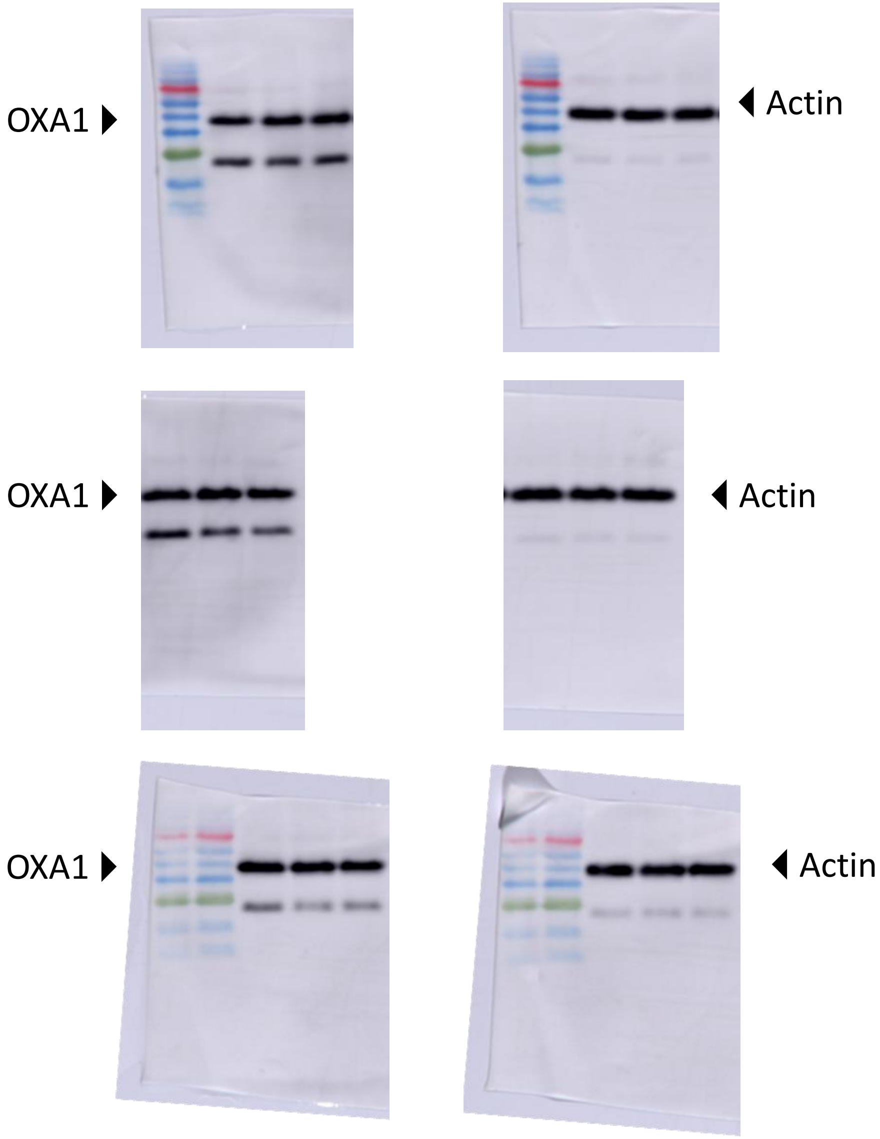
