## Supplemental raw images 2 for "Biochemical and neurophysiological effects of deficiency of the mitochondrial import protein TIMM50"

### TIMM50

(Running order is Untreated / pLL3.7 control / Scr control / Sh1 / Sh2 / Sh3)

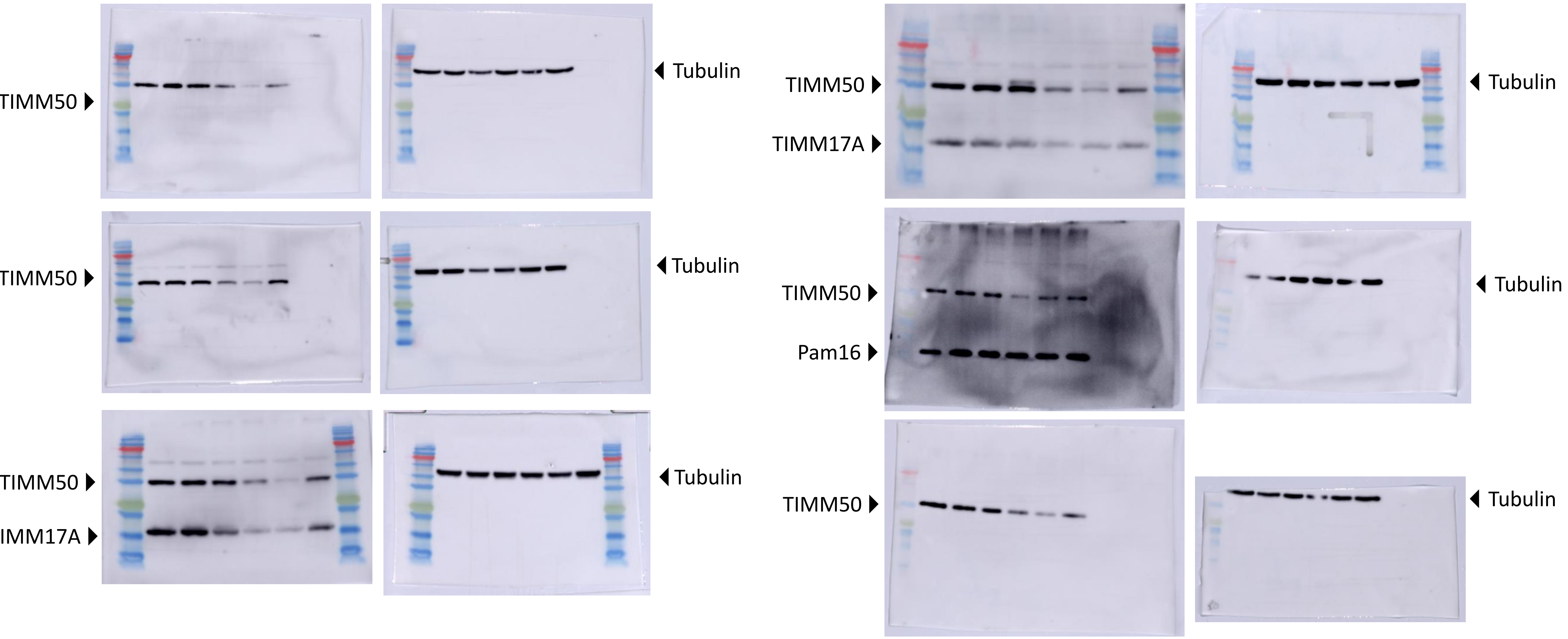

### TIMM50

(Running order is Untreated / pLL3.7 control / Scr control / Sh1 / Sh2 / Sh3)

*The bottom blot has only Scr control and Sh2 samples from 4 separate cultures and a different running order than the rest

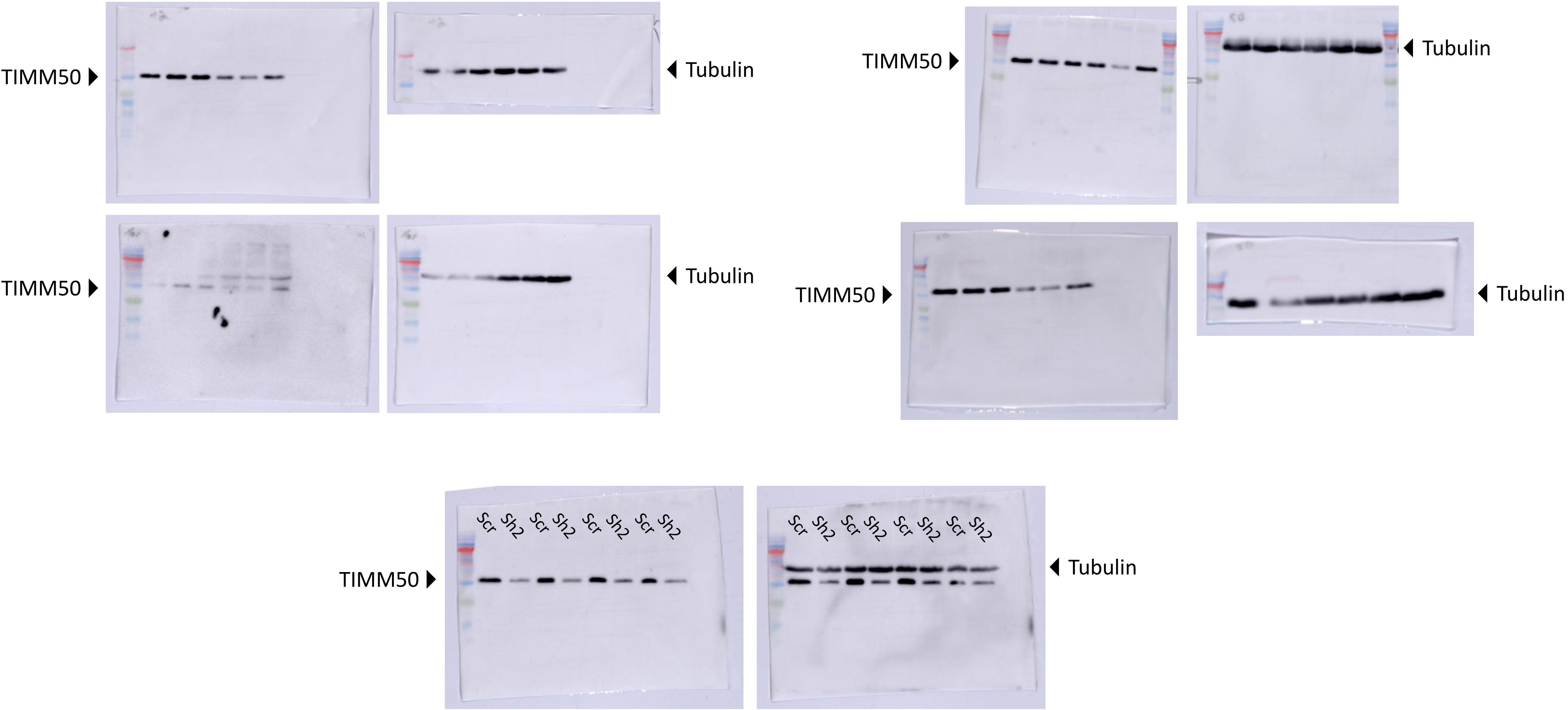

### TIMM23

(Running order is Untreated / pLL3.7 control / Scr control / Sh1 / Sh2 / Sh3)

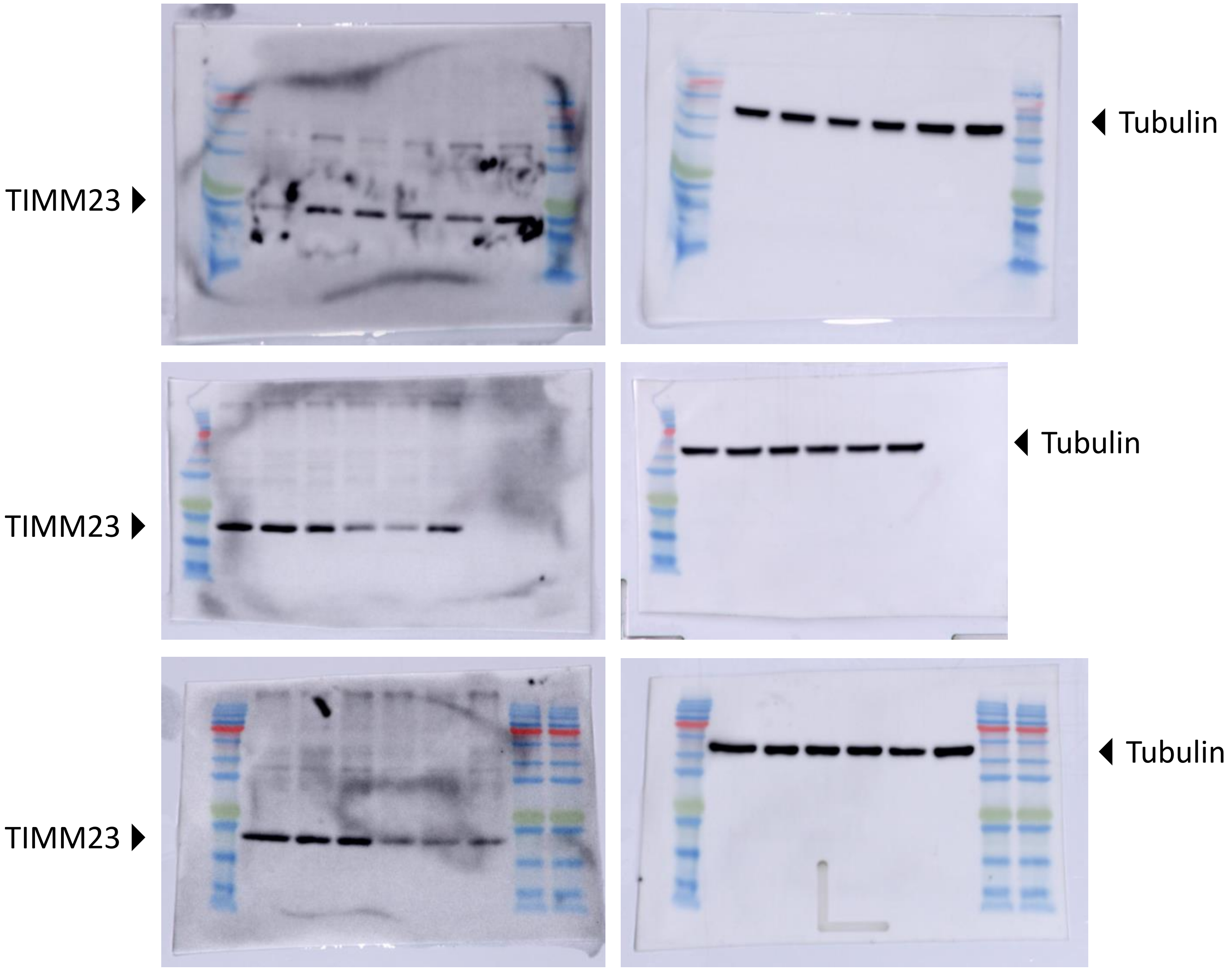

### TIMM17A

(Running order is Untreated / pLL3.7 control / Scr control / Sh1 / Sh2 / Sh3)

*Note that the blot on the right had a different running order

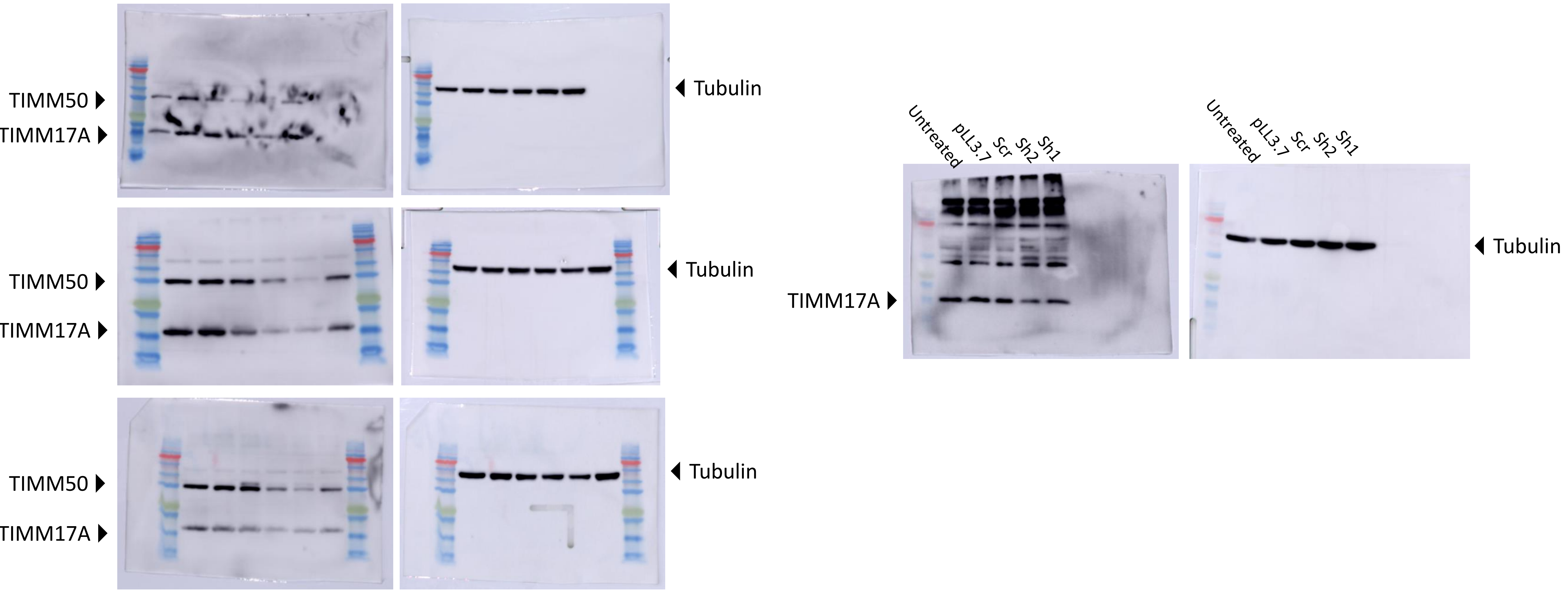

### TIMM17B

(Running order is Untreated / pLL3.7 control / Scr control / Sh1 / Sh2 / Sh3)

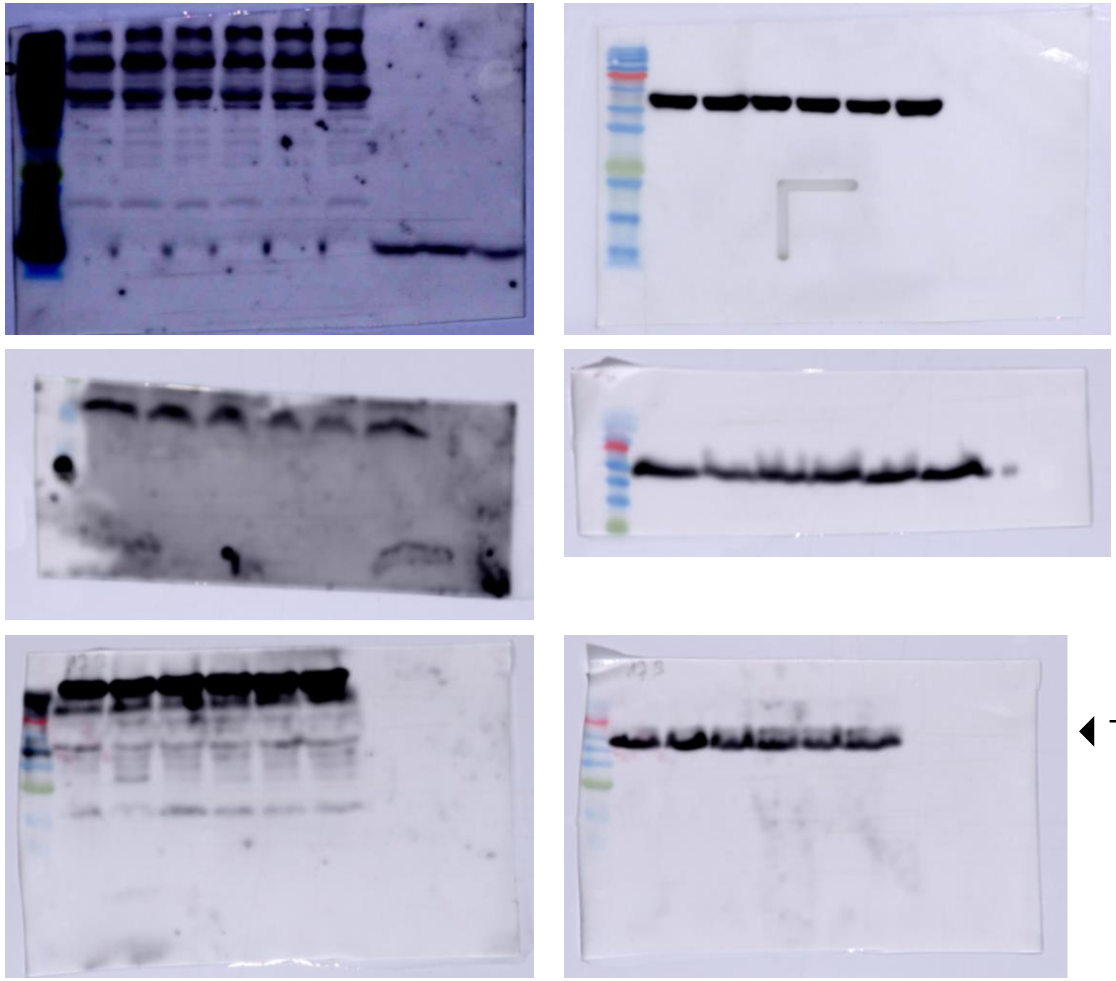

TIMM17B

Tubulin

TIMM17B (Upper part of the membrane was used for Timm21)

Tubulin

TIMM17B

Tubulin

### TIMM21

(Running order is Untreated / pLL3.7 control / Scr control / Sh1 / Sh2 / Sh3)

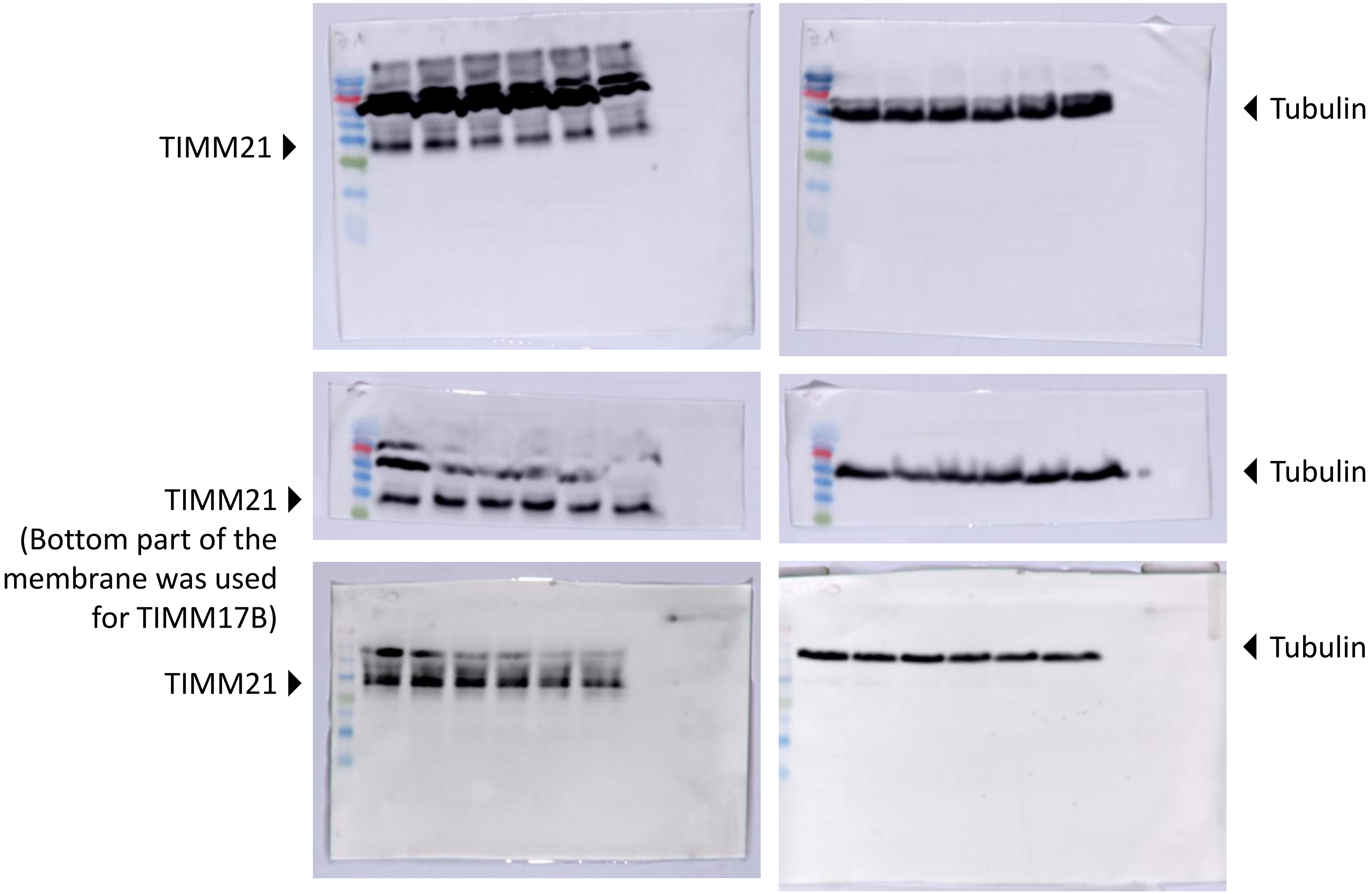

### TIMM44

(Running order is Untreated / pLL3.7 control / Scr control / Sh1 / Sh2 / Sh3)

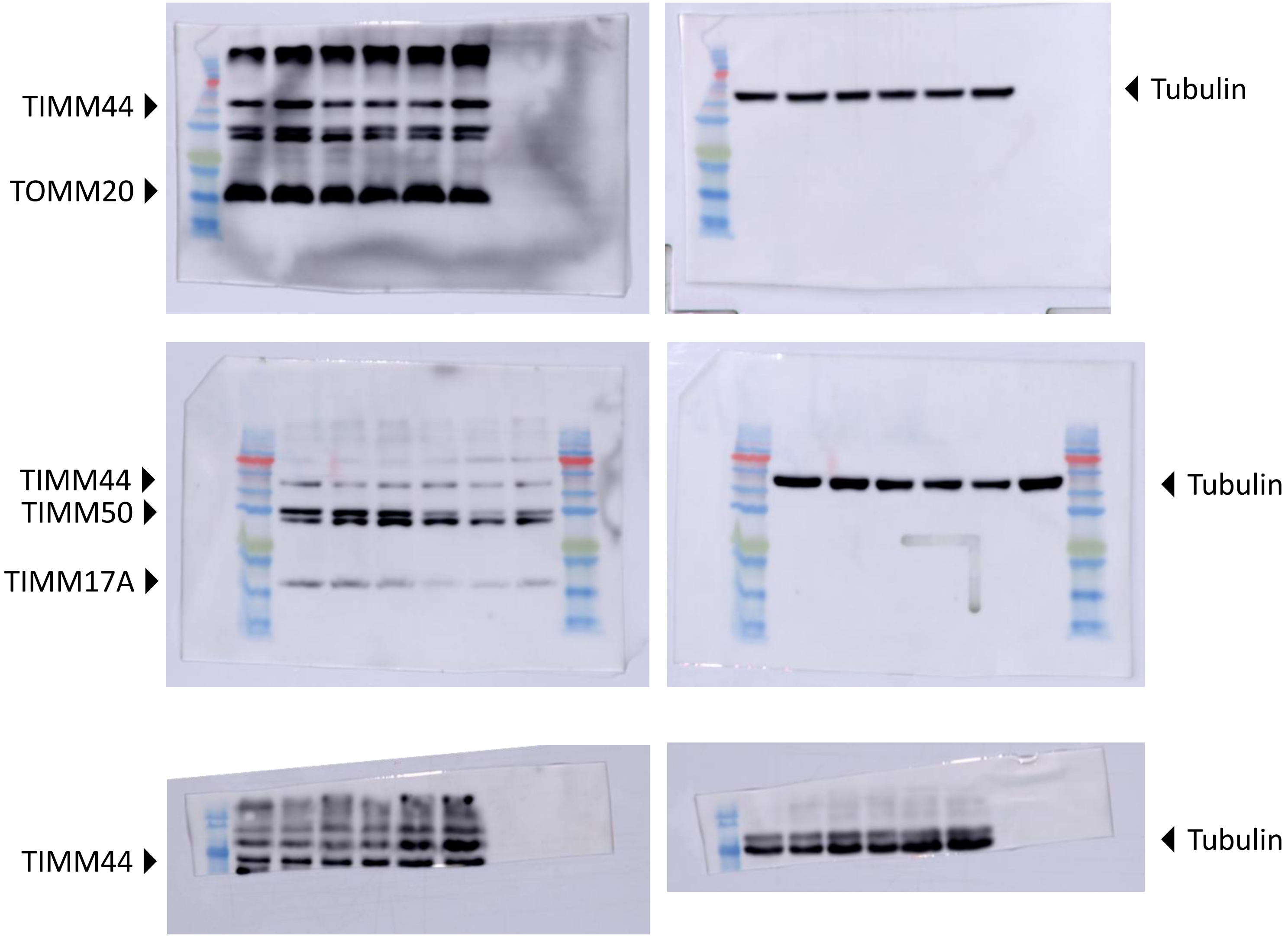

### Pam16

(Running order is Untreated / pLL3.7 control / Scr control / Sh1 / Sh2 / Sh3)

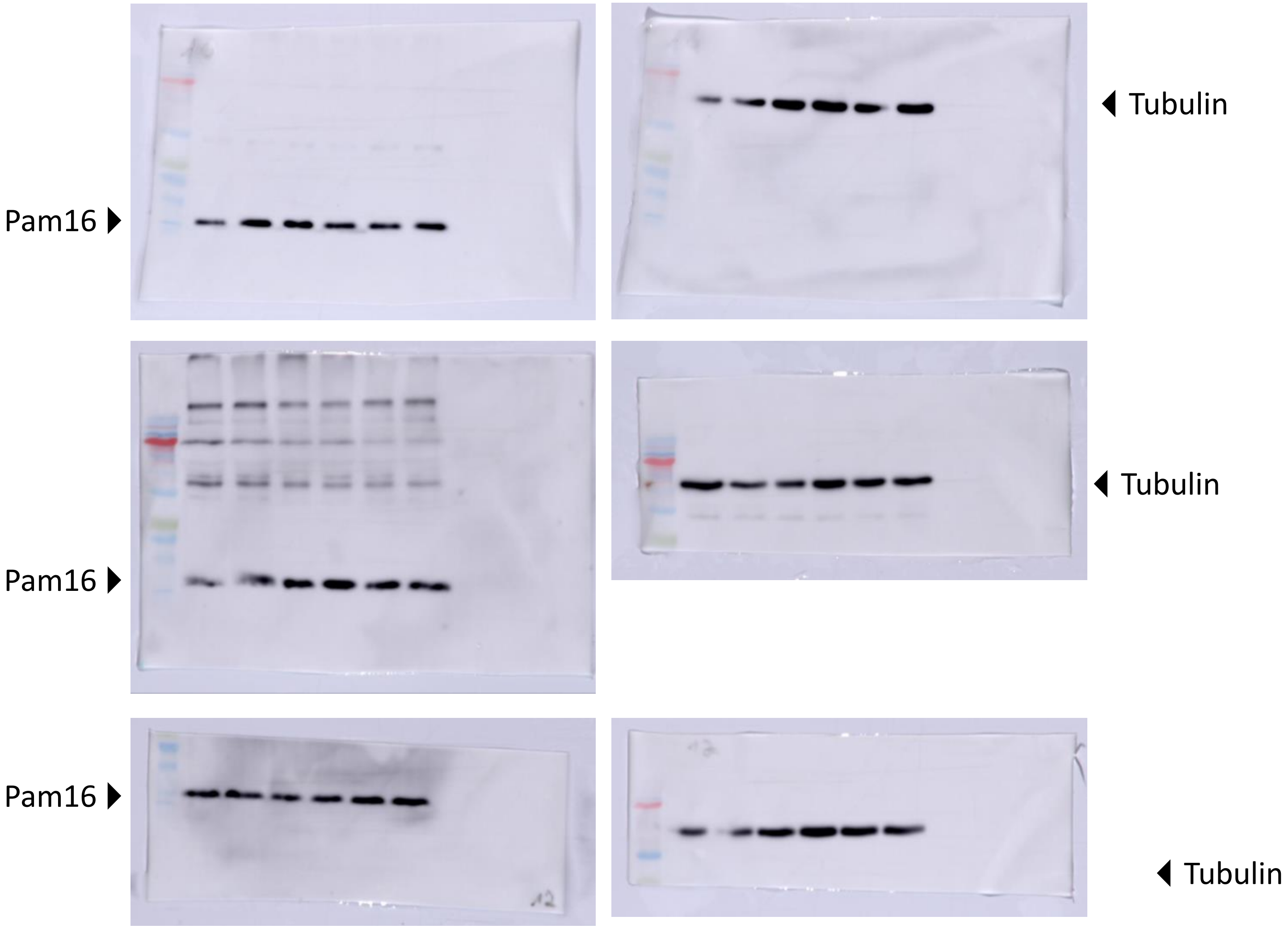

### TOMM40

(Running order is Untreated / pLL3.7 control / Scr control / Sh1 / Sh2 / Sh3)

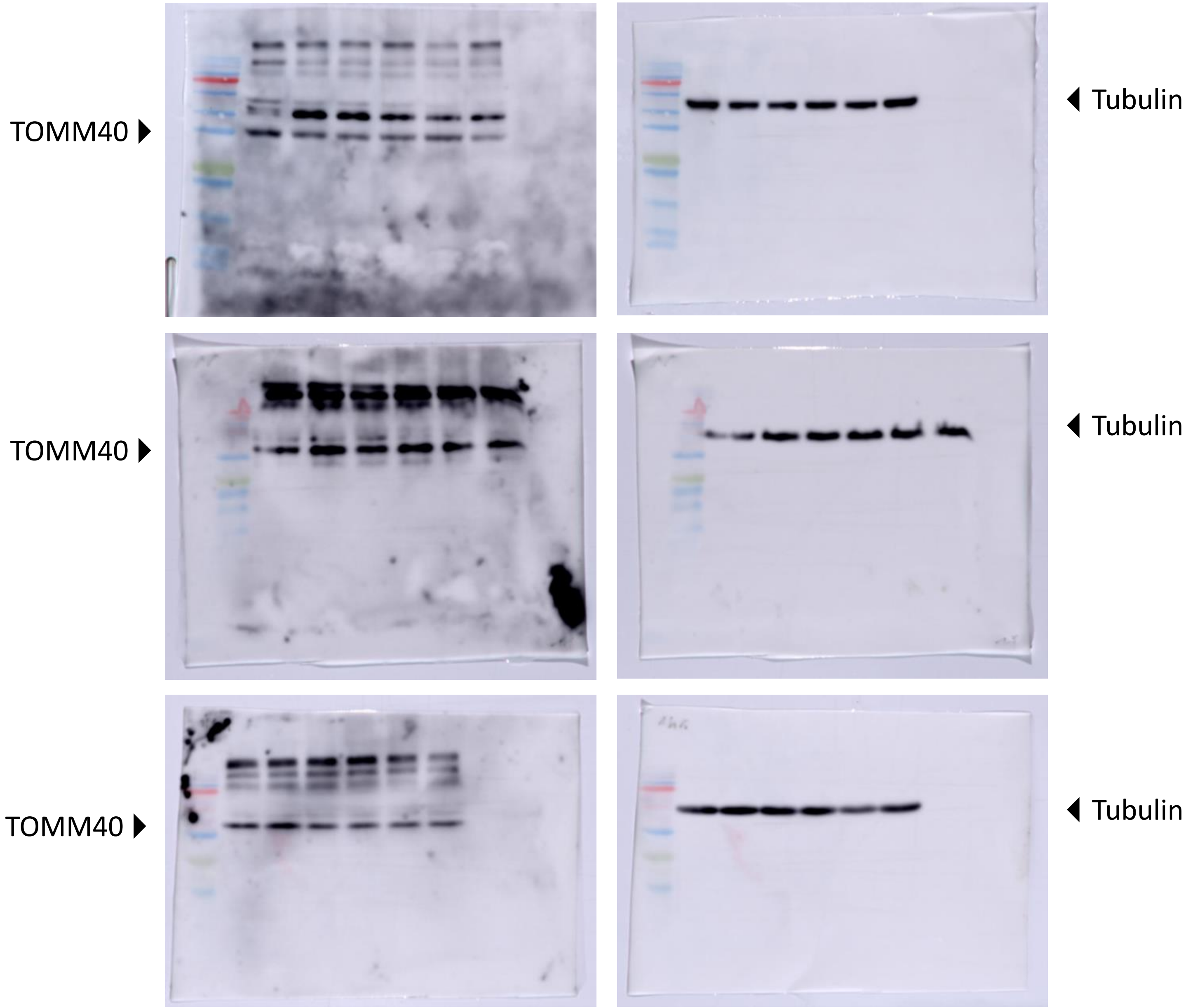

### TOMM20

(Running order is Untreated / pLL3.7 control / Scr control / Sh1 / Sh2 / Sh3)

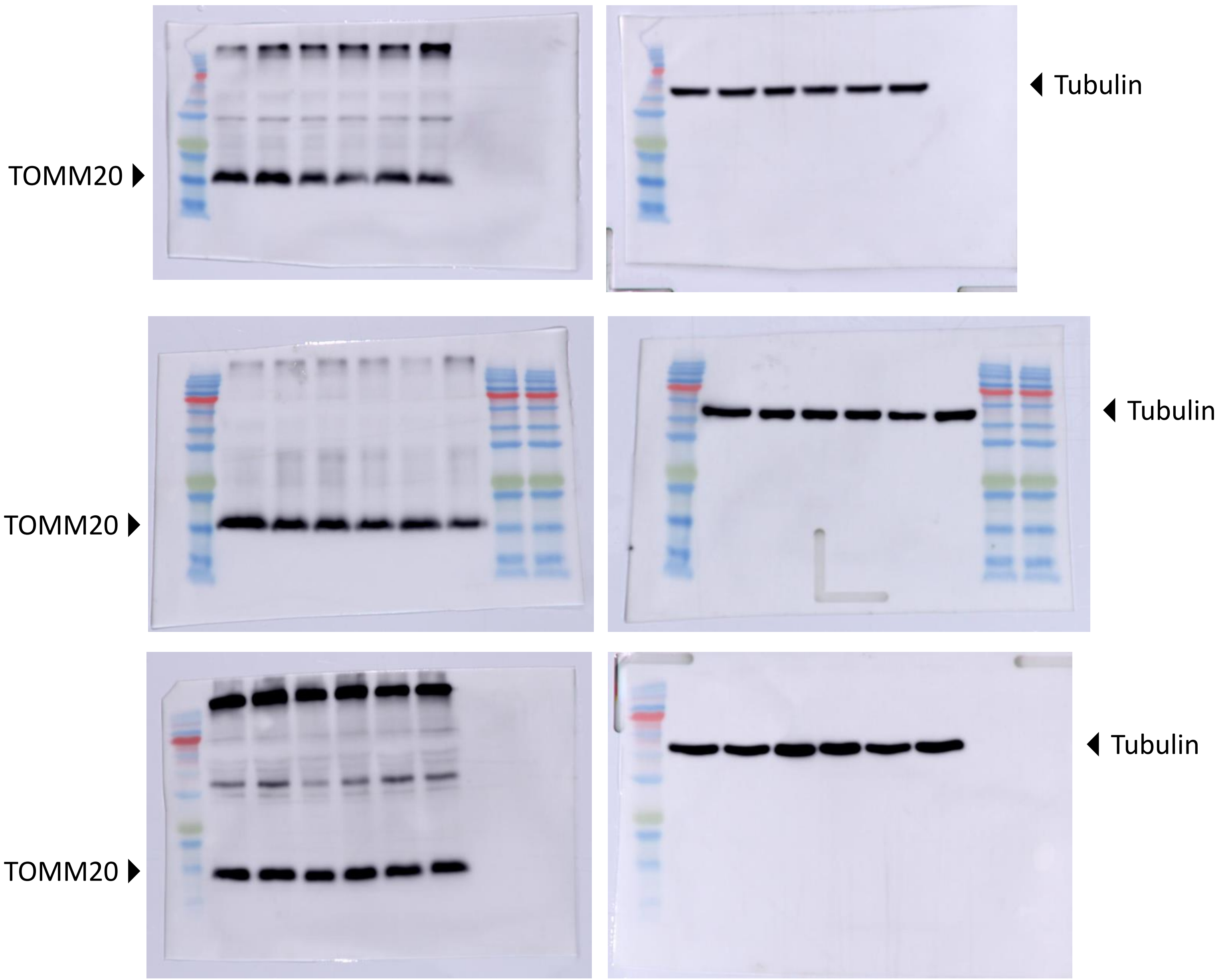

### mtHsp60

(Running order is Untreated / pLL3.7 control / Scr control / Sh1 / Sh2 / Sh3)

mtHsp60 ▶

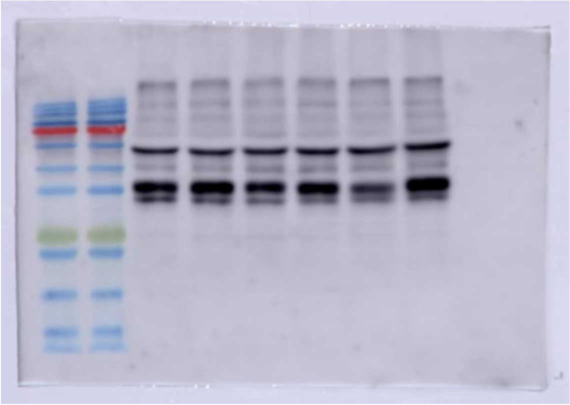

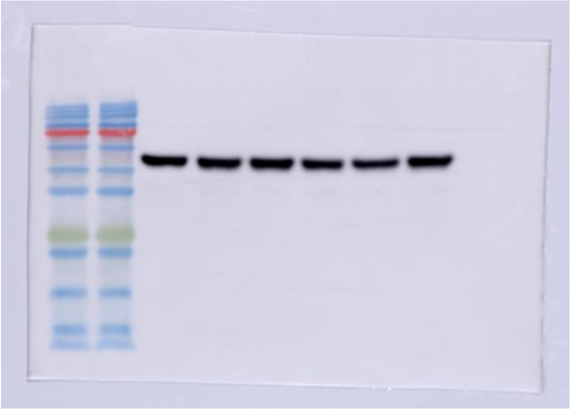

◀ Tubulin

Aconitase-2 ▶
mtHsp60 ▶

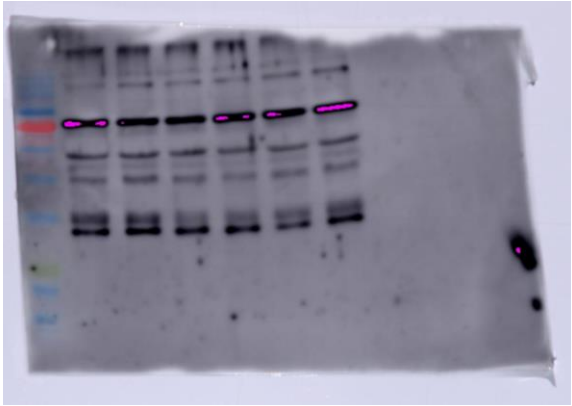

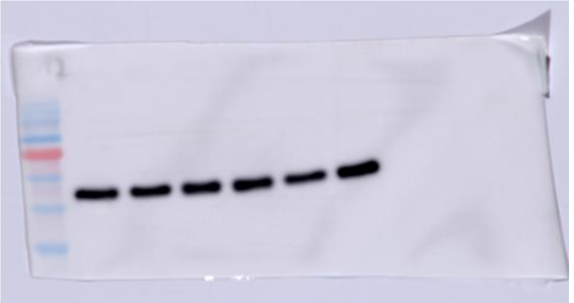

◀ Tubulin

mtHsp60 ▶
(Same membrane was used for Pam16 so the bottom was removed)

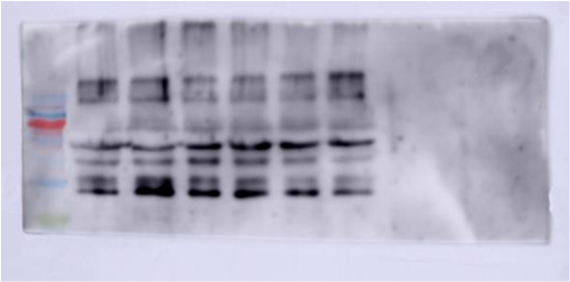

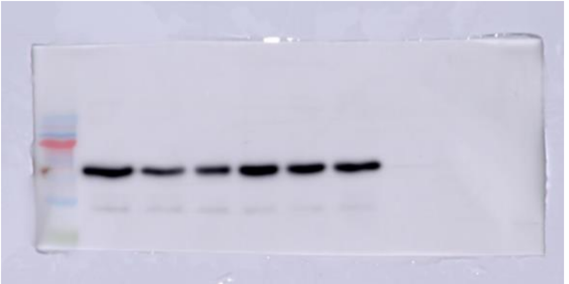

◀ Tubulin

### Aconitase-2

(Running order is Untreated / pLL3.7 control / Scr control / Sh1 / Sh2 / Sh3)

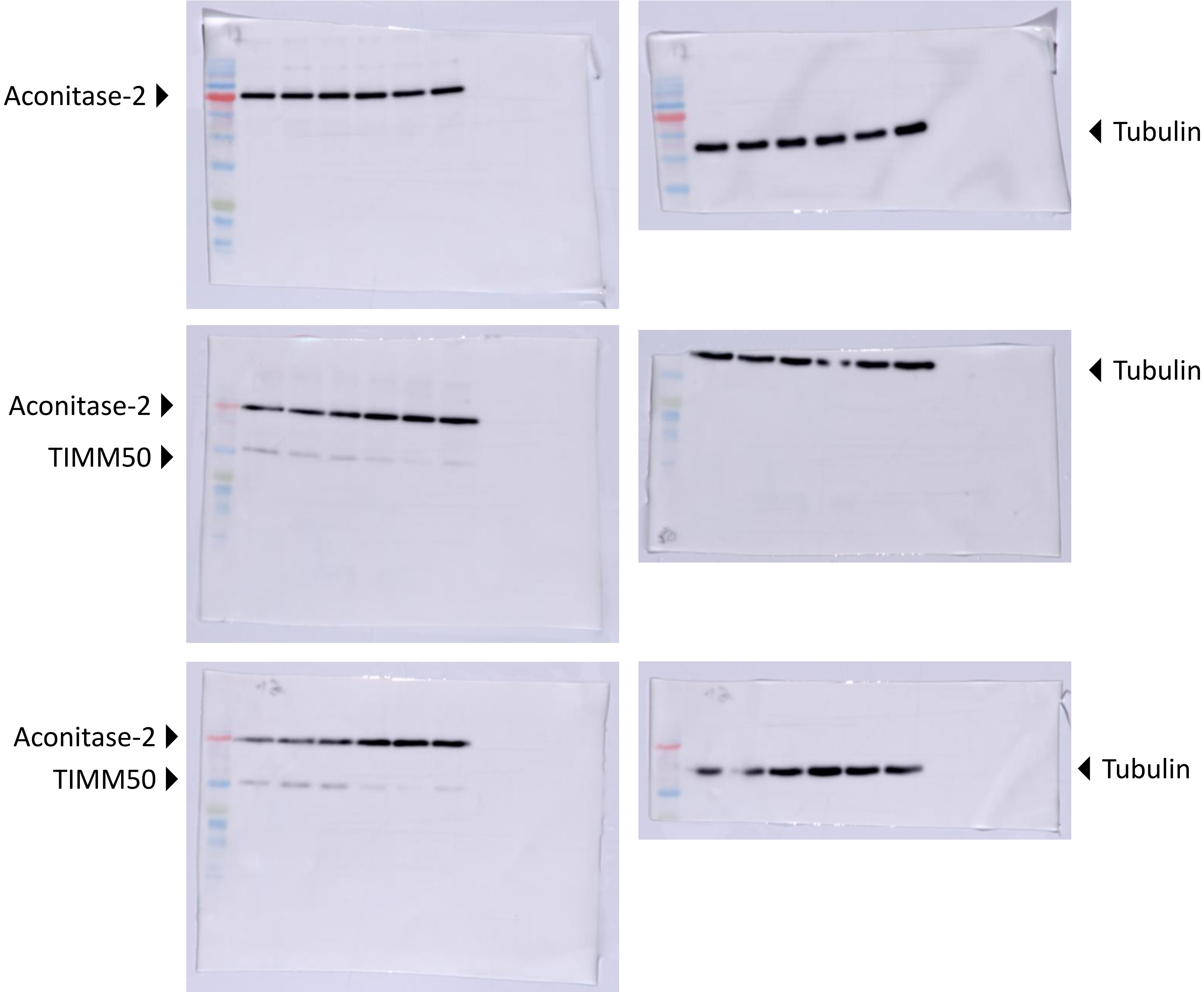

### KCNA2

(Running order is Untreated / pLL3.7 control / Scr control / Sh1 / Sh2 / Sh3)

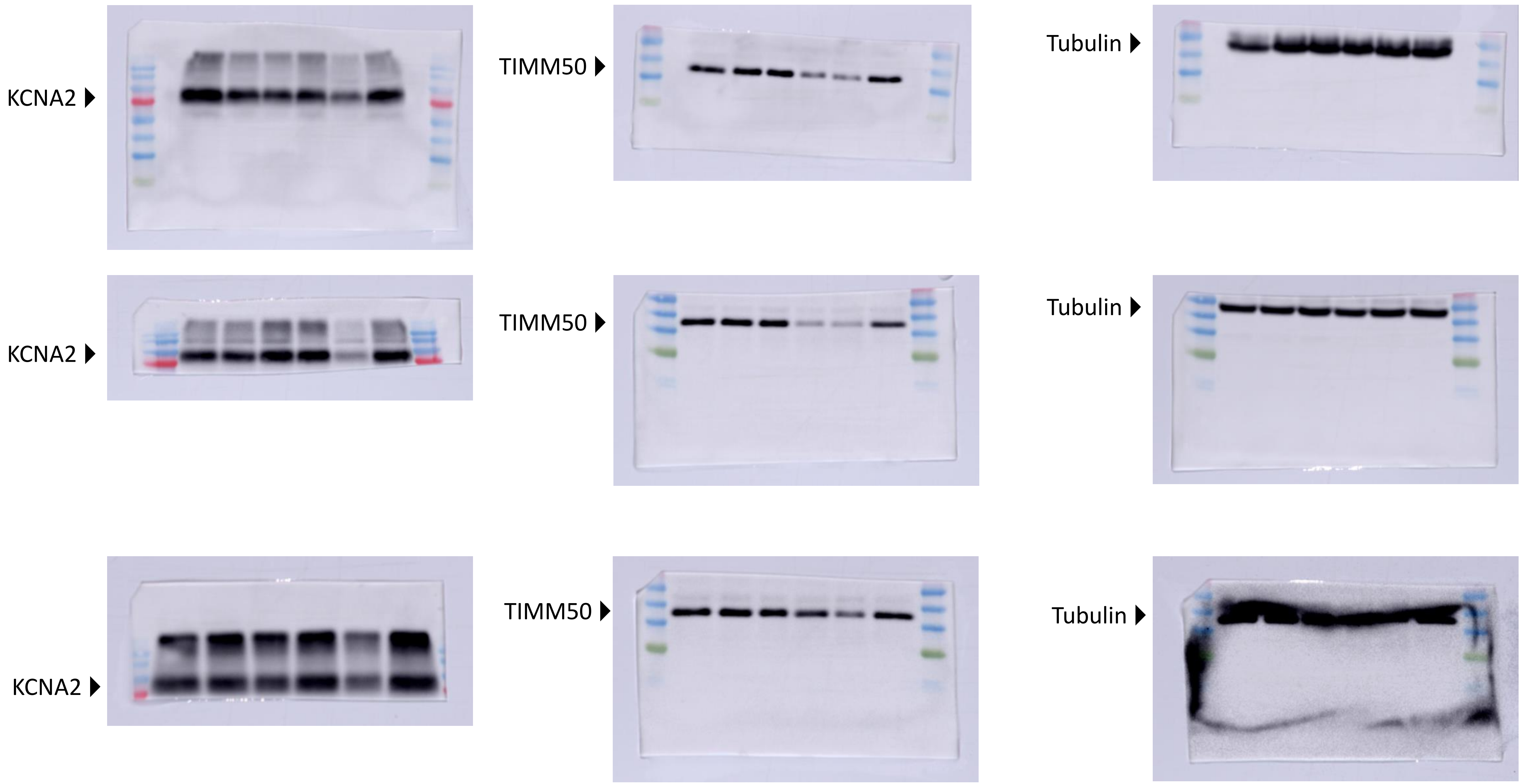
